# Crosskingdom growth benefits of fungus-derived phytohormones in Choy Sum

**DOI:** 10.1101/2020.02.04.933770

**Authors:** Keyu Gu, Shruti Pavagadhi, Yoon Ting Yeap, Cheng-Yen Chen, Sanjay Swarup, Naweed I. Naqvi

**Affiliations:** Temasek Life Sciences Laboratory, 1 Research Link, Singapore 117604; Department of Biological Sciences, 16 Science Drive 4, National University of Singapore, Singapore 117558; Singapore Centre for Environmental Life Sciences Engineering, 60 Nanyang Drive, Singapore 637551; NUS Environmental Research Institute, National University of Singapore, Singapore 117411

**Keywords:** Choy Sum, *Penicillium citrinum*, growth promotion, indole acetic acid, Gibberellin, Cytokinin, Phytohormone

## Abstract

Soil-borne beneficial microbes establish symbioses with plant hosts, and play key roles during growth and development therein. In this study, fungal strains FLP7 and B9 were isolated from the rhizosphere microbiome associated with Choy Sum (*Brassica rapa* var. *parachinen-sis*) and barley (*Hordeum vulgare*), respectively. Sequence analyses of the internal transcribed spacer and 18S ribosomal RNA genes combined with colony and conidial morphology identified FLP7 and B9 to be isolates of *Penicillium citrinum.* Plant-fungus interaction assays revealed that B9, but not FLP7, showed significant growth promotion effect in Choy Sum cultivated in normal soil, whereas FLP7 enhanced Choy Sum growth under phosphate-limiting condition. In comparison to the mock control, B9-inoculated plants showed a 34% increase in growth in aerial parts, and an 85% upsurge in the fresh weight of roots when cultivated in sterilized soil. The dry biomass of inoculated Choy Sum increased by 39% and 74% for the shoots and roots, respectively. Root colonization assays showed that *P. citrinum* associates directly with the root surface but does not enter/invade the roots of inoculated Choy Sum plants. Preliminary results also indicated that *P. citrinum* can promote growth in Choy Sum via volatile metabolites too. Interestingly, we detected relatively higher amounts of indole acetic acid and cytokinins in axenic *P. citrinum* culture filtrate through liquid-chromatography mass-spectrometry analyses. This could plausibly explain the overall growth promotion in Choy Sum. Furthermore, the phenotypic growth defects associated with the Arabidopsis *ga1* mutant could be chemically complemented by the exogenous application of *P. citrinum* culture filtrate, which also showed accumulation of fungus-derived active gibberellins. Our study underscores the importance of trans-kingdom beneficial effects of such mycobiome-derived phytohormone-like metabolites in host plant growth.

## INTRODUCTION

Mycorrhizal microbes play an important role in growth, reproduction, and stress tolerance in plant hosts. The strategies include the production of phytohormones, development of lateral root branching and root hair, and improved absorption of nutrients. For example, plant growth-promoting rhizobacteria (PGPR) colonize roots and enhance plant growth directly and indirectly (Vacheron et al., 2013;Mabood et al., 2014). *Penicillium oxalicum* P4 and *Aspergillus niger* P85 can solubilise phosphate (P) and promote maize growth (Yin et al., 2015). Further studies verified that 7 and 4 organic acids showed strong increase associated with isolate P4 and P85 (Yin et al., 2015). Gibberellin mimics produced by fungi play a vital role in plant growth and development. *Penicillium commune* KNU5379 produces more active variants of Gibberellic acid (GA) such as GA3, GA4 and GA7 (Choi et al., 2005). On the other hand, fungus-derived indole acetic acid (IAA) also plays a significant role in plant growth. For instance, *Penicillium menonorum* displayed growth-promoting activity through IAA and siderophore production (Babu et al., 2015). When inoculated with *Penicillium menonorum* KNU-3, the dry biomass of cucumber roots and shoots increased by 57% and 52%, respectively (Babu et al., 2015). Similarly, in Arabidopsis, three fungal endophytes from water mint can increase the fresh and dry weight of Arabidopsis at 14 and 21 days post inoculation. Among them, *Phoma macrostoma* can increase both root area and depth at 21 days (Dovana et al., 2015). Other isolates from *Phoma* and *Penicillium* showed similar effects. The *Phoma glomerata* and *Penicillium* species. can significantly increase chlorophyll content in leaves, and fresh and dry weight of shoots (Waqas et al., 2012). Further analyses detected active Gibberellins such as GA1, GA3, GA4, and GA7; and auxin in the pure cultures from those two strains (Waqas et al., 2012). *Penicillium pinophilum* formed arbuscular mycorrhizae, which increase the plant dry weight, nitrogen content, P content and photosynthesis rate by 31%, 47%, 57% and 71%, respectively (Fan et al., 2008). *Talaromyces pinophilus*, an endophytic fungus isolated from halophytic plants of Korea can increase the plant height in comparison with the uninoculated wild type (Khalmuratova et al., 2015).

Green leafy vegetables are an important source containing many nutrients. As a non-mycorrhizal Brassicaceae species, Choy Sum is a common vegetable in our daily diet. However, current studies of mycorrhizal microbes mainly focus on bacterial communities, the vegetable associated mycorrhizal fungal species are scarce. In order to identify the phytohormone-secreting (GA, cytokinin and IAA) fungi, which can promote overall growth and biomass increase in green leafy vegetables, we isolated several mycobiome species from the roots of Choy Sum (*Brassica rapa* var. *parachinensis*) and barley (*Hordeum vulgare*). Among the one hundred isolated fungi, two isolates, FLP7 and B9, showed promising growth phenotypes in Choy Sum in the laboratory and in soil-based greenhouse cultivation; through secreted phytohormones such as GA, cytokinin or IAA. Our results demonstrate that symbioses with beneficial fungi play an important role in promoting plant growth and increasing agricultural productivity.

## RESULTS AND DISCUSSION

### Isolation of Beneficial Fungi That Enhance Plant Growth

A fungal strain, termed B9, was isolated from the roots of two-week old barley seedlings. The sequencing results of internal transcribed spacer (ITS), large subunit (LSU) and small subunit (SSU) of 18S nuclear ribosomal RNA genes identified B9 to be a *Penicillium citrinum* isolate. The plant-fungus interaction assays were conducted to check if B9 can induce or promote growth in green leafy vegetables. The results indicated that B9 can significantly increase the growth in Choy Sum plants in sterilized soil as well as in non-autoclaved soil (**Figure 1**). Overall, the B9-inoculated plants were larger and grew taller than the un-inoculated controls (**Figure 1A-1D**). Compared with mock control, the fresh and dry weight of aerial parts in the B9-inoculated plants increased by 34.8%, 39.5%, 41.2% and 25.4% in the sterilized or non-sterile soil, respectively (**Figure 1E-1H**; n=24, *P<0.05*). Similarly, the fresh and dry weight of roots increased by 85.4%, 74.9%, 83% and 42.4% under the respective conditions (**Figure 1E-1H**). These data helped us conclude that the B9 isolate of *P. citrinum* can significantly promote growth in Choy Sum both in the presence or absence of resident commensal microbiota in the rhizosphere.

**Figure 1:**
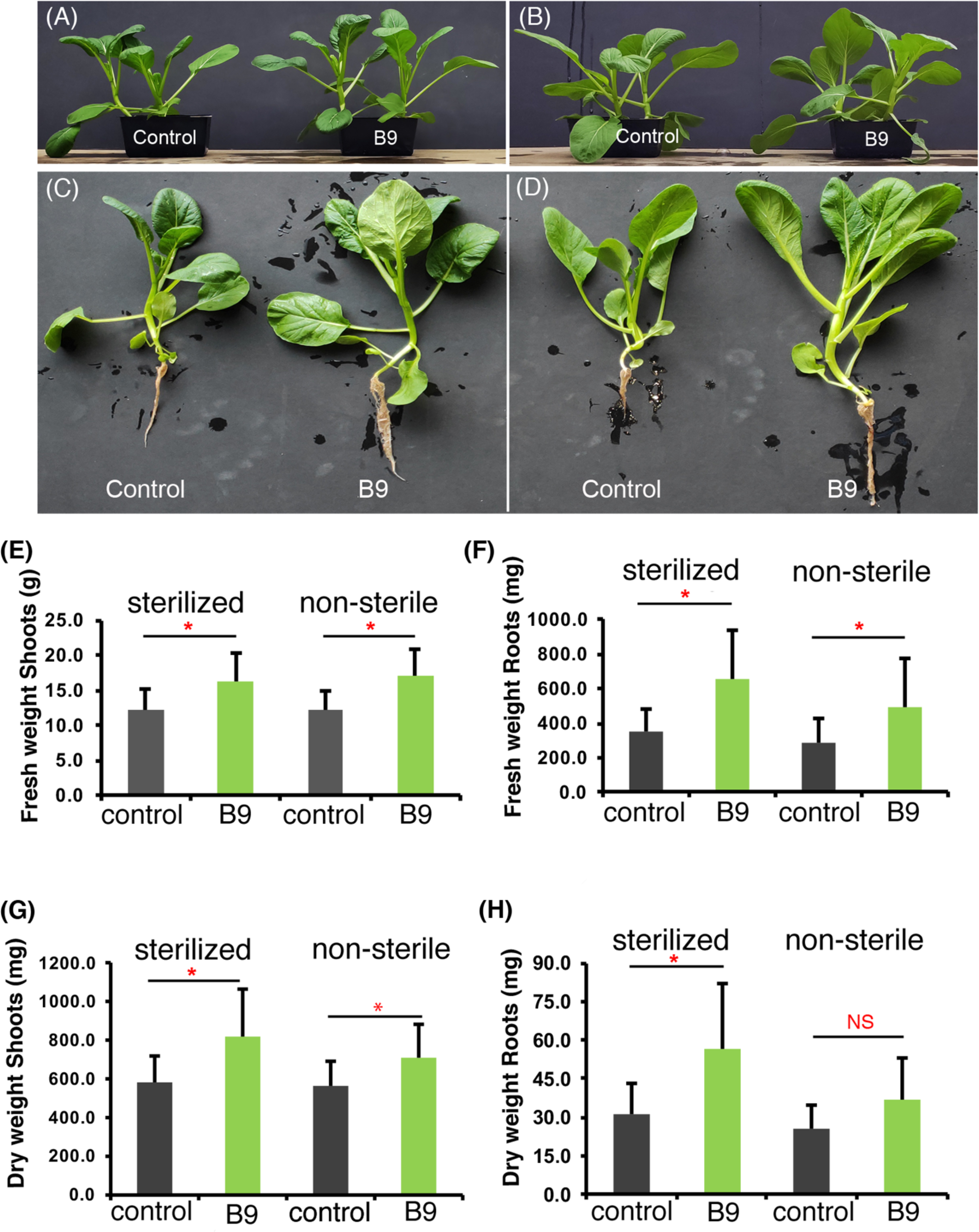
The morphological traits of Choy Sum inoculated with *P. citrinum*, and its plant growth promotion effect in sterilized soil (**A** and **C**) or non-sterile soil (**B** and **D**). (**A** and **C**) The morphological traits of Choy Sum grown for 21 days in autoclaved soil (left, water; right, B9 inoculated); (**B** and **D**) The morphology and growth characteristics of Choy Sum plants grown in non-autoclaved soil for 21 days (left, water; right, B9 inoculated). (**E** to **H**) Bar charts showing quantification of the fresh and dry weight of shoots/aerial parts (**E** and **G**) and roots (**F** and **H**) under the two growth conditions, respectively. Data represents mean ± SD from 3 replicates consisting of 8 plants in each instance. Differences were considered significant at a probability level of p<0.05 (*) or p<0.01(***).

In another experiment, 32 fungal strains belonging to 11 genera were isolated from the roots of Choy Sum grown on Murashige Skoog agar medium with low amounts of phosphate. Of these promising fungal isolates, FLP7 was also tested for growth promotion effects in the common Brassica vegetable Choy Sum in the greenhouse. The results demonstrated that, unlike B9, FLP7 does not promote growth in Choy Sum (**Supplementary Figure S1A-F**). The morphological traits of Choy Sum inoculated with FLP7 were similar to those in the mock controls. Furthermore, the fresh and dry weight of shoots and roots between FLP7 and its corresponding mock control did not show any significant differences (**Supplementary Figure S1C-F**). Interestingly, FLP7 was also identified as *Penicillium citrinum* based on ITS sequence analysis; and showed highly similar phenotypic characteristics as B9 in colony and conidial morphology (**Supplementary Figure S2A-D)**. We conclude that unlike B9, the *P. citrinum* isolate FLP7 has no growth promotion effect in Choy Sum cultivated in soil under such nutrient-rich conditions.

### *P. citrinum* (FLP7) Improves Choy Sum Growth Under Phosphate-limiting Conditions

A previous report showed that the root endophyte *Colletotrichum tofieldiae* (*Ct*) promotes Arabidopsis growth under phosphate deficient condition (Hiruma et al., 2016). However, *Ct* did not display growth promotion under phosphate replete condition (Hiruma et al., 2016). Since FLP7 was isolated from seedlings grown on growth medium containing low levels of phosphate, we tested whether the ability (if any) to promote growth in the host plants is also restricted to phosphate-limiting conditions. To further investigate this, sterilized soil with very low phosphate content (0.11% (w/w)) was used to test this hypothesis. In comparison to mock treatment with sterile water, the Choy Sum seedlings grown on low phosphate soil inoculated with FLP7 conidia showed a significant increase in overall growth and size of the plants (**Figure 2; p<0.05**). The average leaf area, the root length, the dry weight of roots and shoots were 5.7, 2.0, 3.3 and 3.9 times of the control (**Figure 2**). We infer that the FLP7 isolate of *P. citrinum* can indeed improve the overall growth of Choy Sum likely via facilitating the availability and/or uptake of phosphate in the host under low Pi conditions.

**Figure 2:**
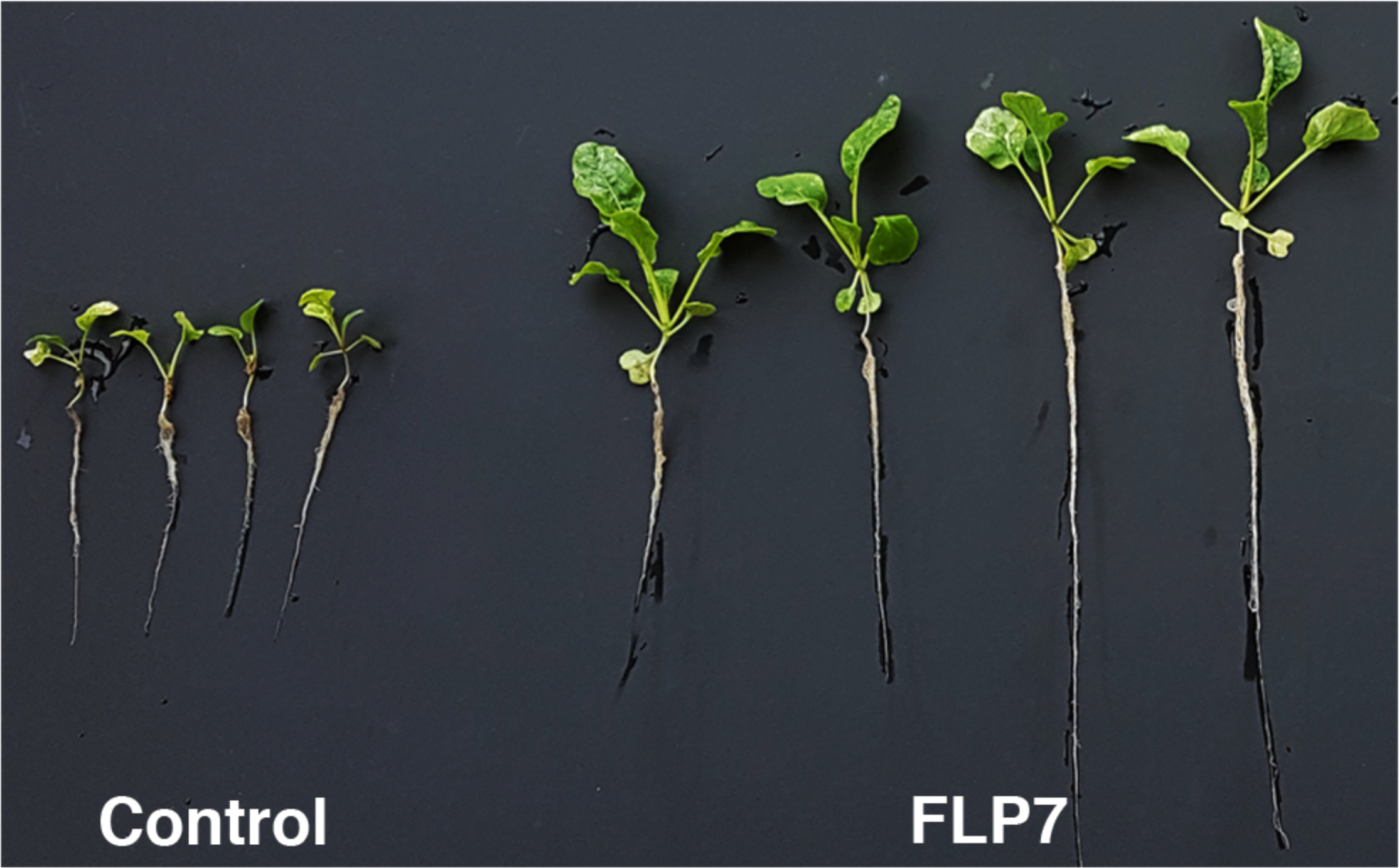
Analyzing the effect of *P. citrinum* FLP7 isolate on the growth of Choy Sum under phosphate-limiting conditions. (**A**) The morphological traits for Choy Sum treated with FLP7 conidia or with water as control. The average leaf area (cm^2^), root length (cm), the dry weight of roots and aerial parts (mg) were determined from both FLP7-treated seedlings and the mock control plants. Data represents mean±SD from 3 replicates consisting of 8 plants in each instance.

### *P. citrinum* Isolates Enhance Choy Sum Growth Via Volatile Secondary Metabolites

Previous studies demonstrated that some beneficial fungi can secrete volatile organic compounds/metabolites to trigger plant growth and development (Hung et al., 2013; Naznin et al., 2013; Jalali et al., 2017). To investigate if FLP7 and B9 can induce similar VOC-based growth stimulation, we incubated the 4-day old Choy Sum seedlings with FLP7 or B9 strain in Phytatray II boxes (Sigma-Aldrich) for 10 days. Barium hydroxide was added to the experimental set-up to quench excess CO_2_, in order to rule out its beneficial effects in plant growth. The tests for such volatile compounds indicated that compared to the mock control (prune agar medium), the size of the seedlings co-cultivated in Phytatray II with B9- or FLP7-isolate was significantly larger (**Figure 3A-C; n=18, *P<0.05***). The fresh and dry weight of shoots and roots of seedlings incubated with FLP7 was 1.46, 1.14, 2.22 and 2.28 times higher than that of the respective mock controls (**Figure 3C-F**). Furthermore, the fresh and dry weight of shoots and roots of the seedlings incubated with B9 was 1.98, 1.63, 2.28 and 2.35 times than that of the un-inoculated control plants (**Figure 3C-F**). We infer that *P. citrinum* secretes some putative volatile compound(s), which are likely responsible for indirectly imparting such beneficial effects on the growth of Choy Sum.

**Figure 3.**
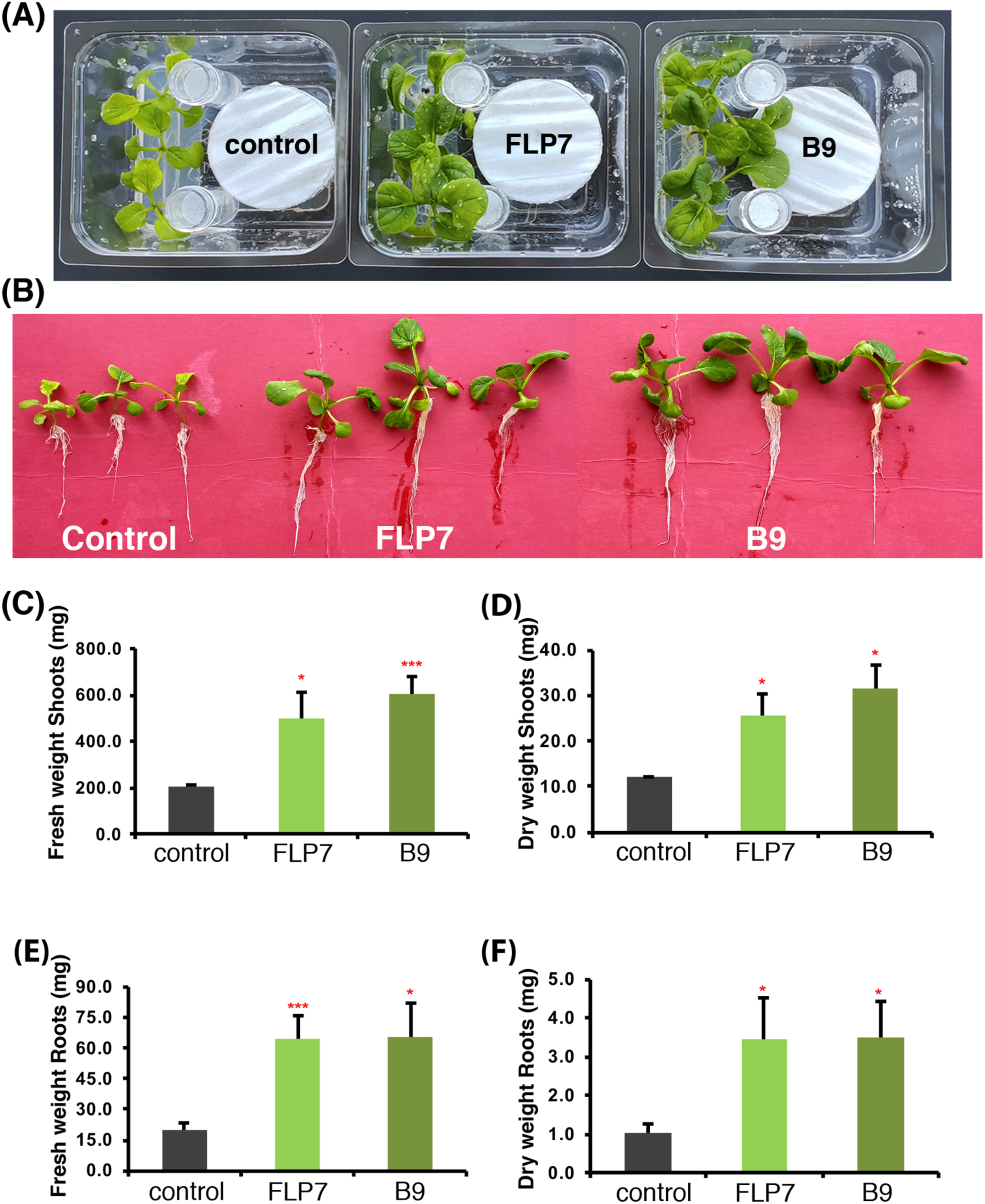
Volatile organic compounds from *P. citrinum* isolates stimulate robust seedling growth in Choy Sum. (**A**) The morphological traits of Choy Sum seedlings grown in tripartite Phytatray II for 10 days; (**B**) the seedlings from (**A**); (**C-F**), quantification of the fresh and dry weight of shoots (**C** and **D**) and roots (**E** and **F**) from the seedlings incubated individually with FLP7 or B9. Data represents mean ± SD from 3 replicates each consisting of 8 plants. Barium hydroxide was used to quench excess carbon dioxide produced during fungal growth or metabolism. Control (mock) inoculation used the growth medium (PA, Prune Agar) without any fungus. Differences were deemed significant at a probability level of p<0.05 (*) or p<0.01(***).

### Analysing the Colonization of Plant Roots by *P. citrinum* Isolates

To further investigate the mode of interaction of FLP7 and B9 in colonizing the roots of Choy Sum, these two isolates were transformed with the gene expressing a cytosolic enhanced green fluorescent protein. The resultant transformants were verified by PCR and sequencing, and the fungal strains expressing the cytosolic eGFP were used for Choy Sum root inoculation assays. The invasive hyphal growth of eGFP-expressing FLP7 was visualized at 12 hours after inoculation. However, no intracellular invasion or colonization within the roots was evident even at 3 days post inoculation (**Figure 4A**). Similarly, the eGFP-tagged B9 strain was used to incubate with 4-day old Choy Sum seedlings. Confocal microscopy revealed that like FLP7, the hyphae of B9 strain of *P. citrinum* also contact and adhere to the surface of Choy Sum roots (**Figure 4B**), but do not enter or colonize the root epidermal cells *per se*. These results demonstrated that the FLP7 and B9 isolates of *P. citrinum* impart the beneficial effects via surface attachment and biotrophic interactions with Choy Sum roots.

**Figure 4.**
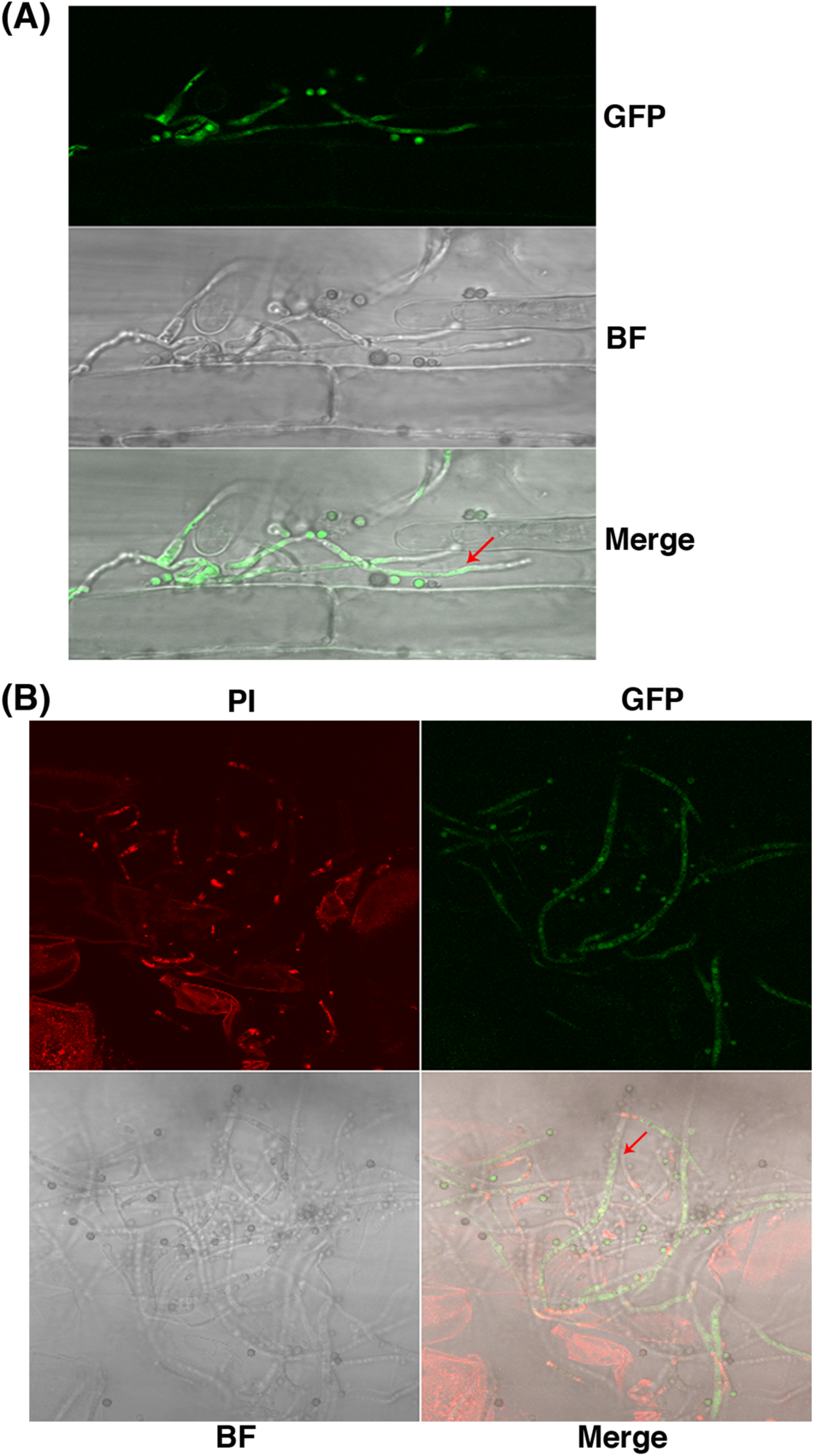
Confocal micrographs of Choy Sum roots incubated with eGFP-expressing FLP7 or B9 strains of *P. citrinum*. Choy Sum roots were incubated with 1 x 10^4^ spores / mL at day 5 post germination. The root colonization was analyzed at 1, 2 and 3 day after incubation. Confocal microscopic images of Choy Sum roots incubated with eGFP-FLP7 (**A**) or B9 (**B**) strains, respectively. Root tissues were stained with Propidium iodide. GFP, green fluorescent protein; BF, bright field, PI, Propidium iodide, Merge, composite of the GFP, PI and BF channels.

### Beneficial *P. citrinum* Isolates Produce Mimics of Gibberellin, Auxin and Cytokinin

Given their strong growth-enhancing effect in the host plants, we decided to evaluate whether *P. citrinum* isolates produce/secrete any growth-promoting secondary metabolites or phytohormones. Towards this end, the axenic culture filtrates of FLP7 and B9 were analysed using liquid chromatography-mass spectrometry (LC-MS) together with the requisite standards for 3 major plant hormones: Gibberellins (GA), Indole acetic acid (IAA/auxin) and Cytokinins. Phytohormone detection was performed using optimized reaction monitoring conditions for the requisite standards as detailed in **Supplementary Figures S3-**S5, and **Supplementary Table S1.** These results demonstrated that the bioactive GAs including GA1 and GA3, and the inactive GA20 variant are present in the culture filtrate of the FLP7 isolate of *P. citrinum* (**Table 1; Supplementary Figure S6A**). However, the results were variable possibly due to the low abundance and/or stability of GAs produced and secreted by the fungus; and the GA-related inter-conversions since only selected GAs were monitored in this study. In order to determine if the GA-like compounds produced by *P. citrinum* are functionally active, an Arabidopsis gibberellin-deficient mutant, *ga1*, and its isogenic wild-type Col-0 accession were germinated on growth medium lacking or containing the culture filtrate of FLP7 or B9 (**Figure 5**). After 9 days, the *ga1* mutant supplemented with FLP7 culture filtrate (but not B9), showed shoot elongation and early flowering (**Figure 5A**). In addition, the wild-type Arabidopsis plants treated with cell-free exudate of FLP7 showed early flowering (**Figure 5B, lower**) compared to the mock treated control (**Figure 5B, upper**). Based on such chemical complementation analysis, we conclude that the *P. citrinum* FLP7 strain indeed produces minor albeit significant amounts of functional GA, which are sufficient *in trans* to suppress the growth defects in the *ga1* mutant of Arabidopsis.

**TABLE 1.**
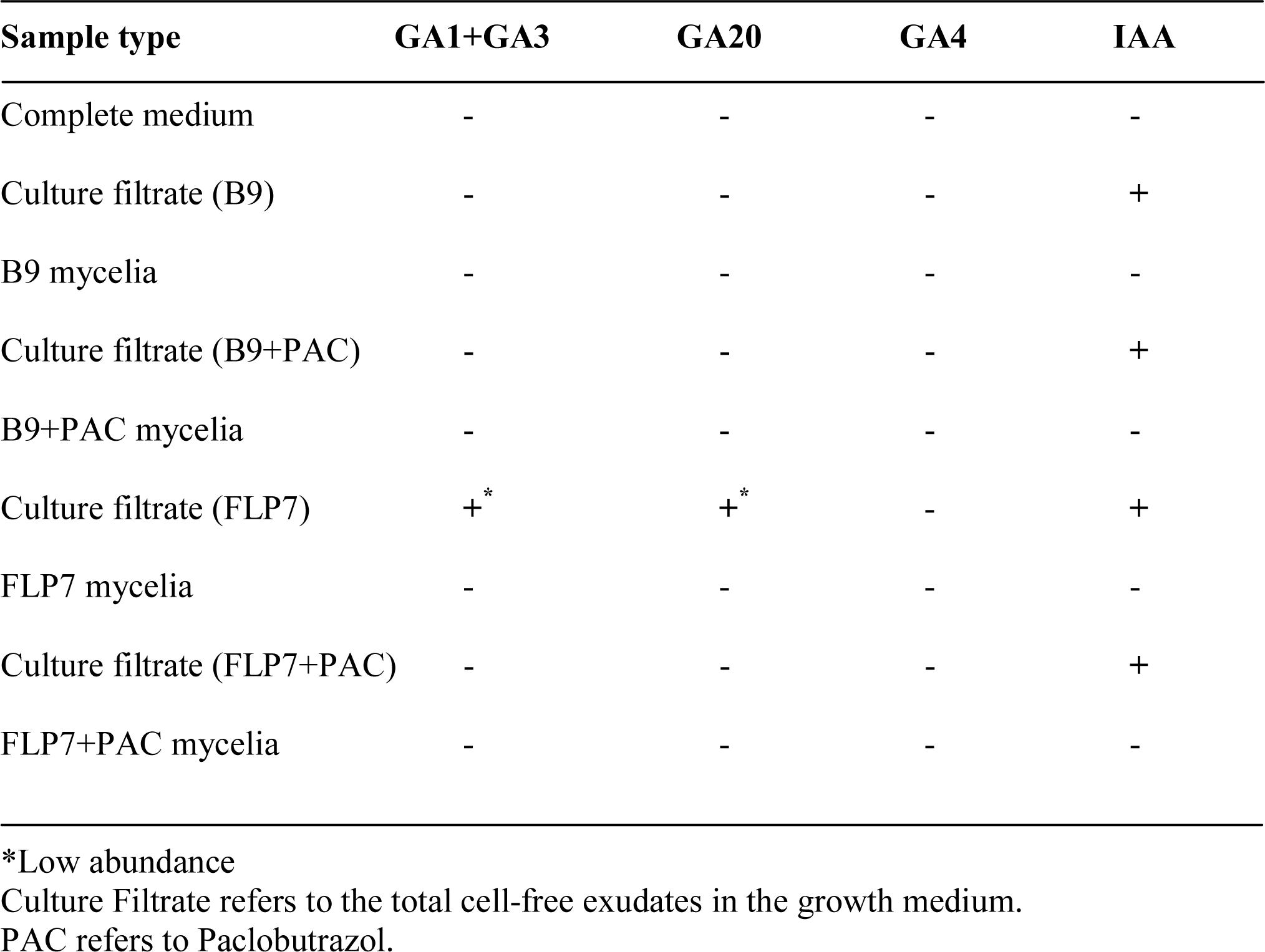
Fungus-derived Gibberellins and IAA produced by *P. citrinum* isolates under various experimental conditions.

| Sample type | GA1+GA3 | GA20 | GA4 | IAA |
| --- | --- | --- | --- | --- |
| Complete medium | - | - | - | - |
| Culture filtrate (B9) | - | - | - | + |
| B9 mycelia | - | - | - | - |
| Culture filtrate (B9+PAC) | - | - | - | + |
| B9+PAC mycelia | - | - | - | - |
| Culture filtrate (FLP7) | + | + | - | + |
| FLP7 mycelia | - | - | - | - |
| Culture filtrate (FLP7+PAC) | - | - | - | + |
| FLP7+PAC mycelia | - | - | - | - |

\*Low abundance
Culture Filtrate refers to the total cell-free exudates in the growth medium.
PAC refers to Paclobutrazol.

**Figure 5.**
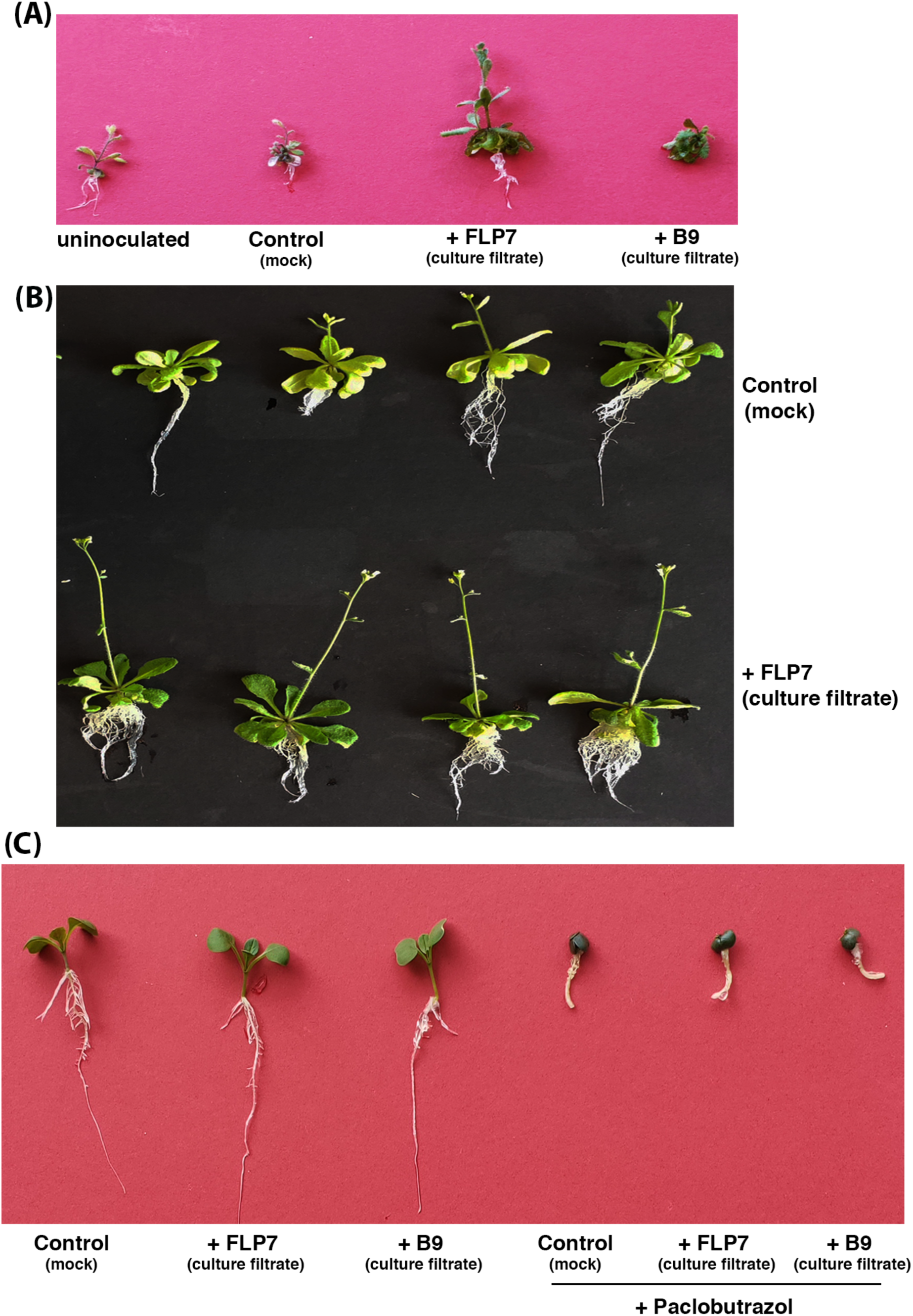
The effect of *P. citrinum* on the growth of Arabidopsis *ga1* mutant and wild-type accession Col-0. (**A**) The morphological traits of Arabidopsis *ga1* mutant in the presence or absence of exudates from *P. citrinum* cultures grown in complete medium. (**B**) Wild-type Arabidopsis Col-0 treated with culture filtrate from *P. citrinum.* Control refers to the mock inoculation with equivalent amount of growth medium without the fungus. (**C**) Gibberellin(s) produced by *P. citrinum* contribute in part to growth promotion in Choy Sum. The culture filtrate from the indicated *P. citrinum* isolate grown in complete medium in the presence or absence of Paclobutrazol (the gibberellin biosynthesis inhibitor) was inoculated on Choy Sum seeds. Control refers to mock inoculation with sterile growth medium in the absence of the fungus.

Likewise, auxin was also found in the culture filtrate of the B9 isolate of *P. citrinum* (**Table 1; Supplementary Figure S6B**). On the contrary, no auxin or gibberellins could be detected from the mock control (un-inoculated growth medium) or in the mycelial extracts (**Table 1**), indicating that the IAA and GA detected in the fungal culture filtrates were most likely secreted by *P. citrinum* and not sourced from the growth medium *per se*. This also suggests that these two fungal phytohormone mimics are likely secreted extracellular metabolites. Based on these results, we conclude that growth benefits imparted by *P. citrinum* are due in part to the secreted fungal phytohormone mimics of gibberellin and/or auxin. We infer that plants could likely receive such auxin derivatives through fungal secretions, and a direct contact or interaction with the roots was likely required for such transkingdom effects of fungal phytohormones.

Paclobutrazol is a known inhibitor of gibberellin biosynthesis in plants (Fletcher et al., 2000;Verma et al., 2010). Gibberellin was undetectable in culture filtrates and/or the corresponding mycelia from FLP7 treated with 10 µM Paclobutrazol (**Table 1**). This result indicates that the biosynthesis of fungal GA is blocked in Paclobutrazol-treated *P. citrinum*, whereas auxin accumulation is unaffected. In contrast, the control set without the inhibitor, showed the presence of gibberellin and auxin in the culture filtrates of both the isolates of *P. citrinum* (**Table 1**).

Next, we investigated the effect of *P. citrinum* exudates on seed germination, growth and development of Choy Sum. As indicated in **Figure 5C**, Choy Sum seeds germinated better and produced robust roots on growth medium supplemented with the culture filtrate from FLP7 or B9. However, the addition of Paclobutrazol-treated growth medium significantly inhibited the germination and growth of Choy Sum seedlings **Figure 5C**), which underscores the specific inhibitory effects of Paclobutrazol on production and activity of fungal gibberellins, and their importance in plant growth promotion by *P. citrinum*.

To elucidate why both FLP7 and B9 can promote growth via volatile metabolites, but only B9 can do so in the soil, both strains were further inoculated in complete medium and grown at 28°C for 7 days in the dark. The culture filtrates were harvested from the two isolates and freeze-dried samples processed for LC-MS analysis. The resultant data showed that relatively higher amount of auxin (a phytohormone mimic) is likely produced and secreted by the B9 isolate of *P. citrinum* as compared to the FLP7 strain (**Supplementary Figure S6B**). Auxin was undetectable in the un-inoculated complete medium (**Table 1**), or in the mycelia from FLP7 or B9 (**Table 1**). To confirm this analysis, we incorporated 10 µM Paclobutrazol into the complete medium together with fungal strains FLP7 or B9 and the samples were analysed by LC-MS. Likewise, only auxin but no Gibberellin was detected in FLP7 and B9 culture filtrates supplemented with the inhibitor (**Table 1**). Similarly, the aforementioned phytohormones were undetectable/absent in the mycelial extracts from FLP7 and B9 treated with the gibberellin inhibitor (**Table 1**).

Apart from Gibberellin and auxin, we also evaluated the presence of two cytokinins in the cell-free culture filtrates of these 2 isolates as well as in media extracts. We detected both trans-Zeatin and trans-Zeatin riboside in media extracts (albeit minor amounts) as well as in B9 and FLP7 culture filtrates. However, as in the case with auxin, we observed relatively higher amounts of these cytokinins in B9 as compared to the FLP7 exudate (**Supplementary Figure S6C**). Taken together, we infer that *P. citrinum* (B9 isolate) produces relatively higher amounts of auxin and two cytokinins in addition to the active Gibberellin derivatives, which together likely lead to the crosskingdom increase in growth in B9-inoculated Choy Sum plants. However, detailed analytical studies are warranted to obtain further conclusive insights about these phytohormones mimics or derivatives in inducing growth in the host plants.

We conclude that the FLP7 and B9 isolates of *P. citrinum* transiently associate with the host root surface, and root colonization *per se* is likely not mandated for the observed beneficial effects. The fungus-derived phytohormones likely play a key role in promoting robust growth and increased biomass in Choy Sum, an economically important urban vegetable crop in Singapore and Asia.

### Similarity and Differences Between the FLP7 and B9 Isolates of *P. citrinum*

PCR amplification using three pairs of primers for ITS, and Large- or Small-subunit of ribosomal genes (**Table 2**), and the sequence analyses indicated that both FLP7 and B9 are isolates of *P. citrinum*. However, when these fungal strains were grown on rich medium for 3 days, different pigments (based on color change) accumulated at the base of the colony (**Supplementary Figure 2A-2B**). The colonies and conidia produced by B9 and FLP7 were highly similar in size and morphology (**Supplementary Figure 2C-2D**). We infer that these phenotypic differences in the colonies of FLP7 and B9 may reflect the metabolic adaptation to different plant host genotypes/ sources and/or the prevalent growth conditions. Additionally, the B9 mycelia might secrete additional as yet unidentified compounds or secondary metabolites during interaction with the host plants. Lastly, both FLP7 and B9 showed growth promotion effect via volatile compounds, but differed in their ability to enhance growth in phosphate-replete conditions; and in the relative levels of the phytohormones gibberellin, auxin and cytokinin produced *in vitro* during mycelial growth. Global metabolomics analyses and whole genome sequencing will likely be required in the future to address these differences between FLP7 and B9 isolates of *P. citrinum*.

**TABLE 2.**
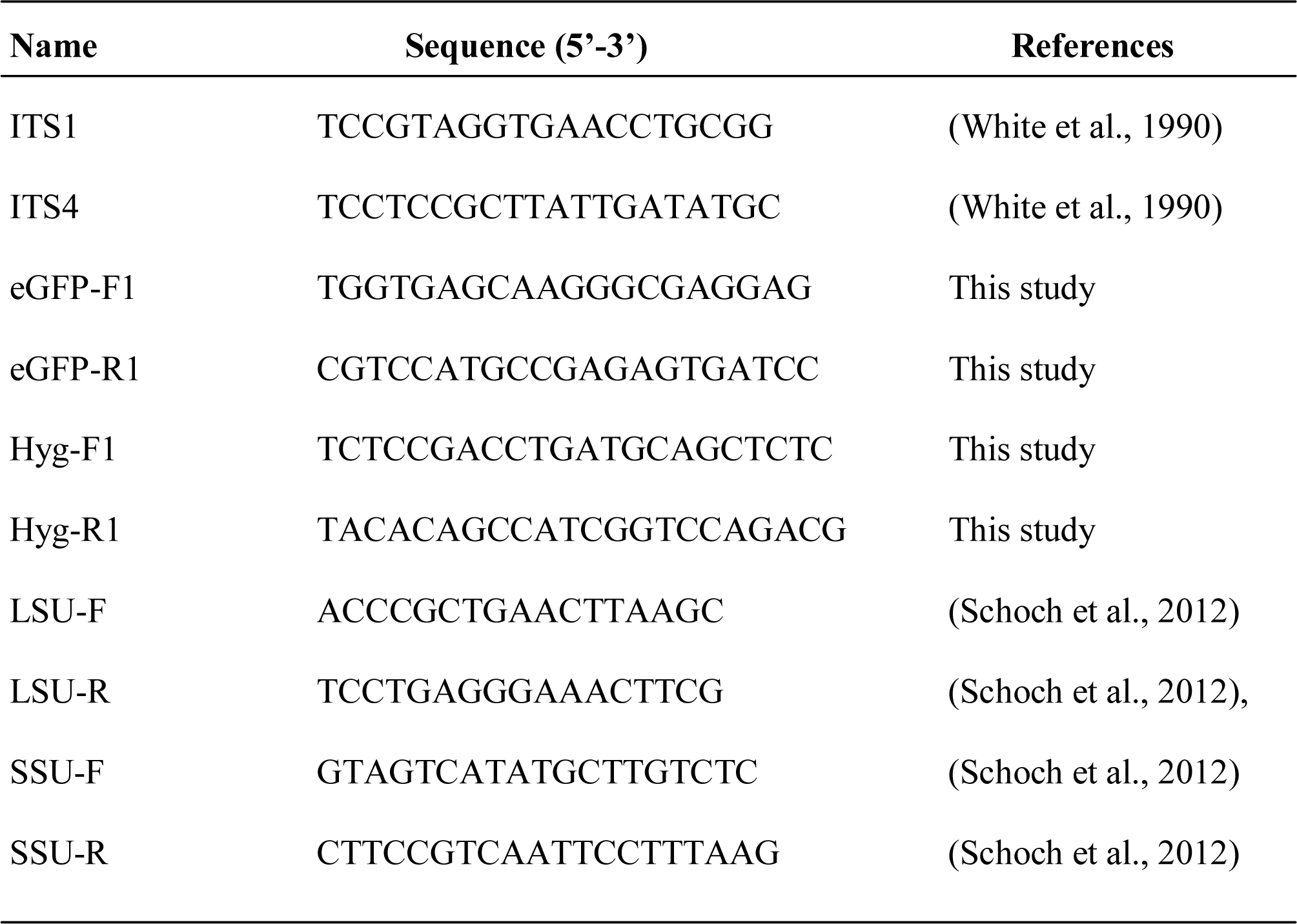
Oligonucleotide primers used in this study.

### Production of Phytohormone Mimics by *P. citrinum*

This study adds to the emerging role and importance of microbiome-derived phytohormone mimics that contribute to the functional aspects of growth benefits in the host plants. Many fungi have been shown to produce phytohormones such as auxin and gibberellins (Hasan, 2002;Choi et al., 2005;Nassar et al., 2005;Rim et al., 2005;Khan et al., 2012;Waqas et al., 2012). For example, auxin can be detected in the cultures of *P. glomerate* and *Penicillium sp*., which increases the plant biomass and related growth parameters under abiotic stress conditions (Waqas et al., 2012). The endophytic yeast *Williopsis saturnus* was found to be capable of producing auxin and indole-3-pyruvic acid *in vitro*, which significantly enhanced the growth in maize plants under gnotobiotic condition and in soils supplemented with or without L-TRP (Nassar et al., 2005). The auxin and Gibberellin(s)produced by *Paecilomyces formosus* were, likewise, implicated in increased growth in rice seedlings (Khan et al., 2012). Similarly, *Fusarium oxysporum*, also produces IAA (Hasan, 2002). In this study, both B9 and FLP7 isolates of *P. citrinum* produced IAA and/or GAs in varying albeit functionally significant amounts during the interaction with Choy Sum **(Table 1)**.

Even though both isolates belong to *P. citrinum*, B9 seems to produce relatively higher amounts of auxin and cytokinin (Supplementary Figure S6), thus indicating the adoption of different strategies or metabolic responses during interaction with the host plants. It remains to be seen whether such phytohormone mimics play a role in fungal growth, development and adaptation *per se*. Future studies will focus on understanding the spatiotemporal regulation of such (fungal) phytohormone mimics and the pathways that regulate the crosskingdom transport to specific subcellular compartments thus leading to enhanced growth in the host tissues.

## CONCLUSION

Under natural conditions, rhizosphere microbes have different interactions with the plant hosts ranging from commensalism to mutualism. Reciprocally, plants can also shape the rhizosphere microbiome for its growth, development, and abiotic- and biotic-stress tolerance. Although many studies have been conducted, the understanding of molecular mechanisms associated with the beneficial microbe and host interactions is still far from complete. Next-generation sequencing technology, combined with the development of metatranscriptomics, metaproteomics and metabolomics, will push forward the understanding between host and the rhizosphere microbes. In this study, we showed that beneficial *P. citrinum* isolates can promote growth in Choy Sum (an important crop for urban farming, and for food and nutritional security) by secreting phytohormone(s) mimics and putative volatile compounds. We demonstrated that rhizosphere fungi can be considered as a useful resource, which can enhance soil fertility and promote plant growth. Integrating the knowledge of mycobiome community composition, beneficial microbial consortia, volatile signals and mutual interactions could aid in sustainable agriculture in an urban setting. Future studies will be directed at understanding the physiology and mechanism-of-action of fungal phytohormone mimics in cross-kingdom growth promotion and resilience in important crop plants.

## MATERIALS AND METHODS

### Fungal Isolation and Identification

Choy Sum seeds were surface sterilized and germinated on Murashige and Skoog medium with low amount of phosphate (12.5 µM). The roots were harvested, ground in 1x phosphate buffered saline and the suspension was diluted and plated on Prune agar medium (40 ml/L prune juice, 1 g/L yeast extract, 2.5 g/L lactose, 2.5 g/L sucrose, 20 g/L agar, pH 6.5) supplemented with Tetracycline (5 µg/ml) + Streptomycin (100 µg/ml) + Carbenicillin (50 µg/ml) + Kanamycin (50 µg/ml). In parallel, the roots were collected from about two-week old barley plants and brought to the laboratory for further processing. The roots were washed with normal water to remove the attached soil and then with distilled water for 30 minutes. After that, the roots were sterilized with 70% ethanol for 1-2 minutes and rinsed with distilled water for 10 minutes. The sterilized roots were cut and ground using a mortar and pestle. The homogenate was filtered with two layers of sterilized miracloth and diluted and plated on growth medium as above. The plates were incubated at 28°C for 2-3 days, and individual fungal colonies were subcultured several times on PA medium. The mycelial mats from a purified single colony were used to colonize 3MM discs, dried and stored at −20°C. The dried mycelial disc was inoculated in 10 ml of CM and incubated at 28°C, 180 rpm for 2 days, and the resulting mycelia collected for DNA extraction.

The genomic DNA from the mycelia was extracted using the MasterPure™ Yeast DNA Purification kit (Lucigen) and used subsequently for PCR amplification. The PCR products were purified and sequenced. The sequences were used for NCBI BLAST analyses. The PCR primer pairs are listed in **Table 2**. The PCR protocol used was 95°C for 5 min; 35 cycles of 95°C for 30 s, 48°C for 30 s, and 72°C for 1.5 min, 72°C for 5 min; 95°C for 5 min, 35 cycles of 95°C for 30 sec, 52°C for 30 sec, and 72°C for 1.5 min, 72°C for 5 min and 95°C for 5 min, 35 cycles of 95°C for 30 sec, 60°C for 30 sec, and 72°C for 30 sec, 72°C for 5 min, respectively.

### Plant Growth-Promotion Assays

Seeds from Choy Sum or the Arabidopsis *ga1* mutant were surface sterilized and placed on Murashige Skoog medium for germination. The Choy Sum seedlings were transplanted to autoclaved or non-autoclaved soil at 4 dpi, respectively. The *ga1* mutant seedlings were transplanted to Phytatray II boxes containing MS medium supplemented with cell-free culture filtrate from FLP7 or B9. The selected fungal mycelial mat grown on PA medium at 28°C for 3 days in darkness, and then cultivated under constant light for 5 days. The conidia were collected and diluted to 1-5 x 10^5^ spores / mL for inoculation. The inoculated plants were placed in a growth chamber for 2 days and then cultivated in greenhouse until 21 dpi. The experiments were repeated three times each using 8-10 seedlings. To assay for growth-promoting volatile metabolites, the Choy Sum seeds were sterilized and grown on MS medium as described above. The four-day old seedlings in triplicate were transferred to Phytatray II boxes with MS medium, together with fungal strains grown on prune agar, and the boxes were incubated at 25°C, 70% relative humidity (RH) (day) and 23°C, 50% RH (night) for 10 days. Barium hydroxide was added to the experimental set-up to quench excess CO_2_, in order to rule out its indirect beneficial effects in plant growth.

### Agrobacterium-Mediated Transformation of FLP7 or B9

Agrobacterium-mediated transformation of target fungi was performed as described previously (Zheng et al., 2015). *Agrobacterium tumefacians* strain AGL1 carrying the appropriate Transfer-DNA vector/plasmid was grown at 28°C in LB medium containing 100 μg/ml Kanamycin overnight. The overnight AGL1 culture was diluted to OD_600_=0.15 in induction medium (10 mL/L K salts (20.5% K_2_HPO_4_, 14.5% KH_2_PO_4_; M salts: 3% MgSO_4_-7H_2_O, 1.5% NaCl), 20 mL/L M salts, (20%) NH_4_NO_3_ 2.5 mL/L, (1%) CaCl_2_ 1 mL/L, (0.01%) FeSO_4_ 10 mL/L, glucose 5 mM/L, MES 40 mM/L, glycerol 0.5%) containing 100 μg/mL kanamycin and 200 μM acetosyringone, and incubated at 28°C with gentle shaking at 160 rpm for 6 h. Simultaneously, conidia (fungal spores) were harvested from fully grown fungal cultures (on prune agar medium under light for about 1 week) and re-suspended to 1×10^6^/mL in distilled water. A sterile 0.45 μM nitrocellulose filter membrane was placed on induction medium containing 200 mM acetosyringone. A mixture of equal volume (100 μl each) of the AGL1 culture and the fungal conidial suspension was spotted and air-dried on the filter membrane. The plate was then incubated at 28°C for 48 hours. After the co-culture, all the growth on the filter membrane was scraped into 2 mL of sterile PBS (containing 200 mM Cefotaxime, 60 mg/mL Streptomycin and 100 mg/mL Ampicillin) and was vortexed briefly. The re-suspension in PBS was plated equally (200 μl) onto ten CM selection medium plates containing 200 μg/mL Cefotaxime (to kill *Agrobacteria*), 60 μg/mL Streptomycin, 100 μg/mL Ampicillin, and 250 μg/mL Hygromycin. The selection plates were incubated at 28°C until the transformed fungal colonies appeared (typically 3-5 days). The individual colonies were selected for mycelium preparation and DNA extraction as above. The primer pairs used for PCR amplification are listed in **Table 2**.

### Fungal Interaction/Colonization Assays In Choy Sum Roots

The Choy Sum seed germination and seedling preparation were as described above. The GFP-expressing FLP7 and B9 strains were prepared as above. The seedlings were submerged in the conidial suspension containing 1 × 10^4^ spores / mL. The interaction between Choy Sum and the GFP-tagged FLP7 or B9 was analyzed by laser-scanning confocal microscopy (Exciter, Zeiss) using the 10x water-immersion and 63x oil objectives. The excitation/emission wavelength (Ex/Em) was 488 nm/505-550 nm.

### Plant Growth Assay Under Low Phosphate (P) Using FLP7 Isolate

The Choy Sum seeds were sterilized and germinated on Murashige Skoog medium with low phosphate (12.5 µM) for 4 days. The germinated seedlings were transplanted to autoclaved rice soil (with 0.11% (w/w) phosphate) and grew for 21 days like above. During the growth, no fertilizer was added. The average leaf area (cm^2^), root length (cm), the dry weight of roots and aerial part (mg) were calculated for both control and FLP7-treated seedlings.

### Extraction And Purification Methods For Detection of Phytohormones in *P. citrinum*

Certified standards of Gibberellins GA1, GA4, GA20 were purchased from OiChemIm Ltd (Czech Republic). Certified GA3 standard, IAA standard and formic acid were purchased from Sigma-Aldrich (USA). Trans-zeatin and trans-zeatin riboside standards were provided by Prakash Kumar (Singapore). Acetonitrile with 0.1% formic acid (Optima LC-MS grade) and methanol (Optima LC-MS grade) were obtained from Fluka Honeywell. Milli-Q water was used for preparation of mobile phase (Millipore, USA). Prime HLB SPE cartridge (200 mg, 6 cc) and Oasis MCX (200 mg, 6 cc) cartridge were supplied by Waters Corporation, UK.

Standard stock solutions (1000 μg/mL) of GA1, GA3, GA4, GA20, Auxin, trans-zeatin and trans-zeatin riboside were prepared in methanol respectively and stored at −20°C in the dark. Stock solutions were used to prepare working standard solutions for analytical experiments. Extraction procedures for Gibberellin and Auxin were adopted from the methods described previously (Khan et al., 2011; Khan et al., 2012; Waqas et al., 2012). Briefly, the liquid complete medium inoculated with FLP7 or B9 was incubated at 28°C at 180 rpm for 7 days. The resulting culture was centrifuged, and the culture filtrate used for LC-MS analysis. Lyophilized fungal culture filtrate was extracted with ethyl acetate containing formic acid and was loaded onto preconditioned solid-phase extraction cartridge. Subsequently, the column was washed with distilled water and the sample was eluted with acidified methanol. The eluate was evaporated to dryness and reconstituted in 50% methanol for further LC-MS/MS analysis.

Extraction of cytokinins (Trans-zeatin and Trans-zeatin riboside) and subsequent sample clean-up and purification were done using methods adapted from Morrison et al. (2015). Briefly, cell-free filtrates were snap-frozen, lyophilized and subsequently homogenized in cold (−20°C) modified Bieleski No. 2 extraction buffer (Methanol: Water: Formic Acid; CH_3_OH:H_2_O:HCO_2_H [15:4:1, v/v/v]). Samples were allowed to extract passively, twice at −20°C and pooled supernatants were dried in a speed vacuum concentrator at ambient temperature (UVS400, Thermo Fisher Scientific, USA). Dried supernatant residues were reconstituted in 1 mL 1M HCO_2_H and subjected to solid phase extraction on a mixed mode, reverse-phase, cation-exchange cartridge (Oasis MCX 6 cc; Waters, UK). Trans-zeatin and trans-zeatin riboside were eluted with 0.35 M NH_4_OH in 60% CH_3_OH. Samples were evaporated and stored at −80°C prior to analyses. Samples were reconstituted in initial mobile phase conditions (95:5 H_2_O:CH_3_OH with 0.08% acetic acid (CH_3_CO_2_H)) prior to analyses.

### Liquid Chromatography–Mass Spectrometry

LC–MS data were acquired on an Agilent 1290 Infinity coupled to an Agilent 6400 series Triple Quadrupole (Agilent, USA). Ultra-high performance liquid chromatography (UHPLC) system was integrated with Agilent 6490 controlled by MassHunter software B.06.00.

For detection of gibberellins and auxin, 10 µL of extracts was chromatographed on a Zorbax RRHD SB-C18 (50 mm length x 2.1 mm diameter, 1.8 µm particle size) (Agilent, US) with column temperature set at 50°C and auto-sampler temperature was set at 4°C. The mobile phase consisted of water acidified with 0.1% formic acid (Solvent A) and acetonitrile acidified with 0.1% formic acid (Solvent B). A gradient elution (flow rate 300 µL/min) consisting of 5% solvent B for 1 min followed by a linear gradient of 100% solvent B at 10.5 min which was maintained till 13.4 min followed by 5% solvent B at 13.5 min to 16.5 min for re-equilibration. Mass spectrometric detection was performed with a Triple Quadrupole in negative mode with an Agilent Jet Stream ESI (G1958-65138) ion source using optimized monitoring reactions (**Supplementary Figure S3-S4; Supplementary Table S1**).

For detection of Trans-zeatin and Trans-zeatin riboside, 10 µL of extracts were chromatographed on Kinetex C18 column (2.6 µm C18 100 Å, 100 x 2.1 mm) (Phenomenex, USA) with column temperature set at 50°C and auto-sampler temperature was set at 4°C. The mobile phase consisted of water acidified with 0.08% acetic acid (Solvent A) and methanol (Solvent B). A gradient elution (flow rate 300 µL/min) was used consisting of 5% of solvent B at 0 min followed 45% solvent B at 4 min; 75% B at 5 min, followed by 95% B at 5.1 and was maintained till 6.1 min followed by 5% solvent B at 6.2 min to 8.2 min for re-equilibration. Mass spectrometric detection was performed with a Triple Quadrupole in positive mode with an Agilent Jet Stream ESI (G1958-65138) ion source using optimized monitoring conditions (**Supplementary Figure S5; Supplementary Table S1**).

The mass spectrometer settings were as follows for all the phytohormones: source temperature 250°C, gas flow 12 L/min, nebulizer gas pressure 35 psi, sheath gas temperature 350°C, sheath gas flow 11 L/min. Data were recorded in the multiple reaction monitoring mode. All data collection, mass spectrometric and statistical analyses were carried out with Mass Hunter Workstation software package: MH Acquisition B.05.00, MH Qualitative Analysis B.06.00. (Agilent Technologies, USA). All samples were randomized before LC-MS analyses.

### Statistical Analysis

The data and comparison with controls were represented by using mean with standard error. The significance of differences between the control and treatments was statistically evaluated by GraphPad (https://www.graphpad.com/quickcalcs/ttest1.cfm). Differences were considered significant at a probability level of p<0.05 (*) or p<0.01(***).

## DATA AVAILABILITY

All datasets for this study are included in the manuscript and the Supplementary Files.

## AUTHOR CONTRIBUTIONS

GK, NN designed the experiments. GK, PS, CC, YT performed the experiments; and GK co-wrote the manuscript with NN. PS helped in metabolite analyses and manuscript revision. GK, SS and NN analysed the data.

## ACKNOWLEDGMENTS

We thank the Fungal Pathobiology group for useful discussions and suggestions. We thank Yu Hao for sharing the Arabidopsis *ga1* mutant; and Prakash Kumar for providing the cytokinin standards. We would also like to acknowledge NUS Environmental Research Institute for technical support.

