## Supplementary Figures S1-S6 for "Crosskingdom growth benefits of fungus-derived phytohormones in Choy Sum"

**Supplementary Figure S1**

(**A** and **B**) The growth characteristics of Choy Sum under mock inoculation trays on the left), or with conidia from *P*. *citrinum* isolate inoculated on the surface ([method (M1)], or with conidia inoculated in close contact with seeds and covered with sterilized soil (method (M2)).
(**C to F**) Bar graphs representing quantification of the fresh and dry weight of shoots and roots from the aforementioned experiments.

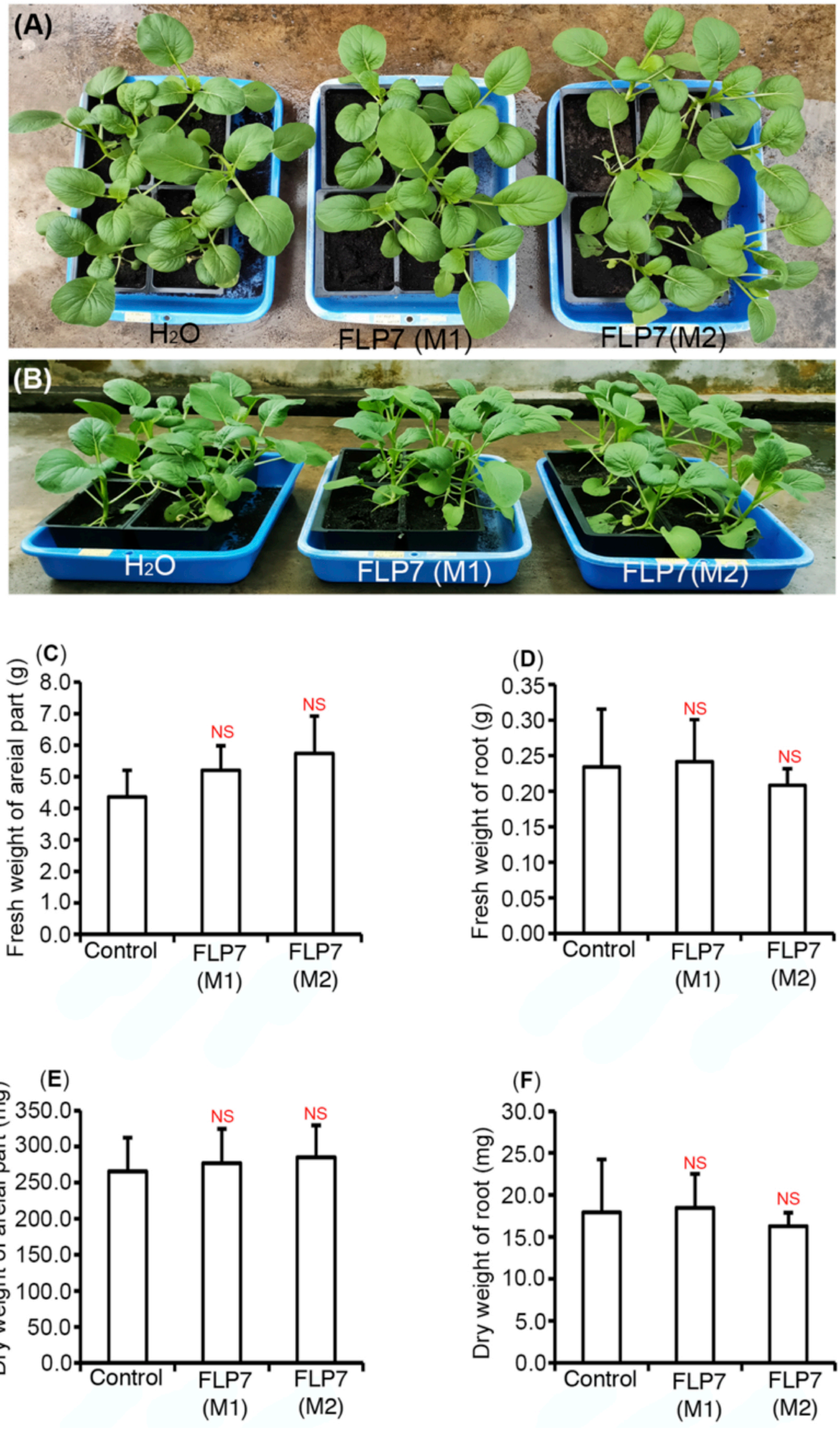

**Supplementary Figure S2**

The morphological characteristics of *P. citrinum* FLP7 and B9 isolates grown on prune agar (**A**) top view, (**B**) bottom view. (**C** and **D**) Morphology of conidia from the indicated isolates of *P. citrinum*.

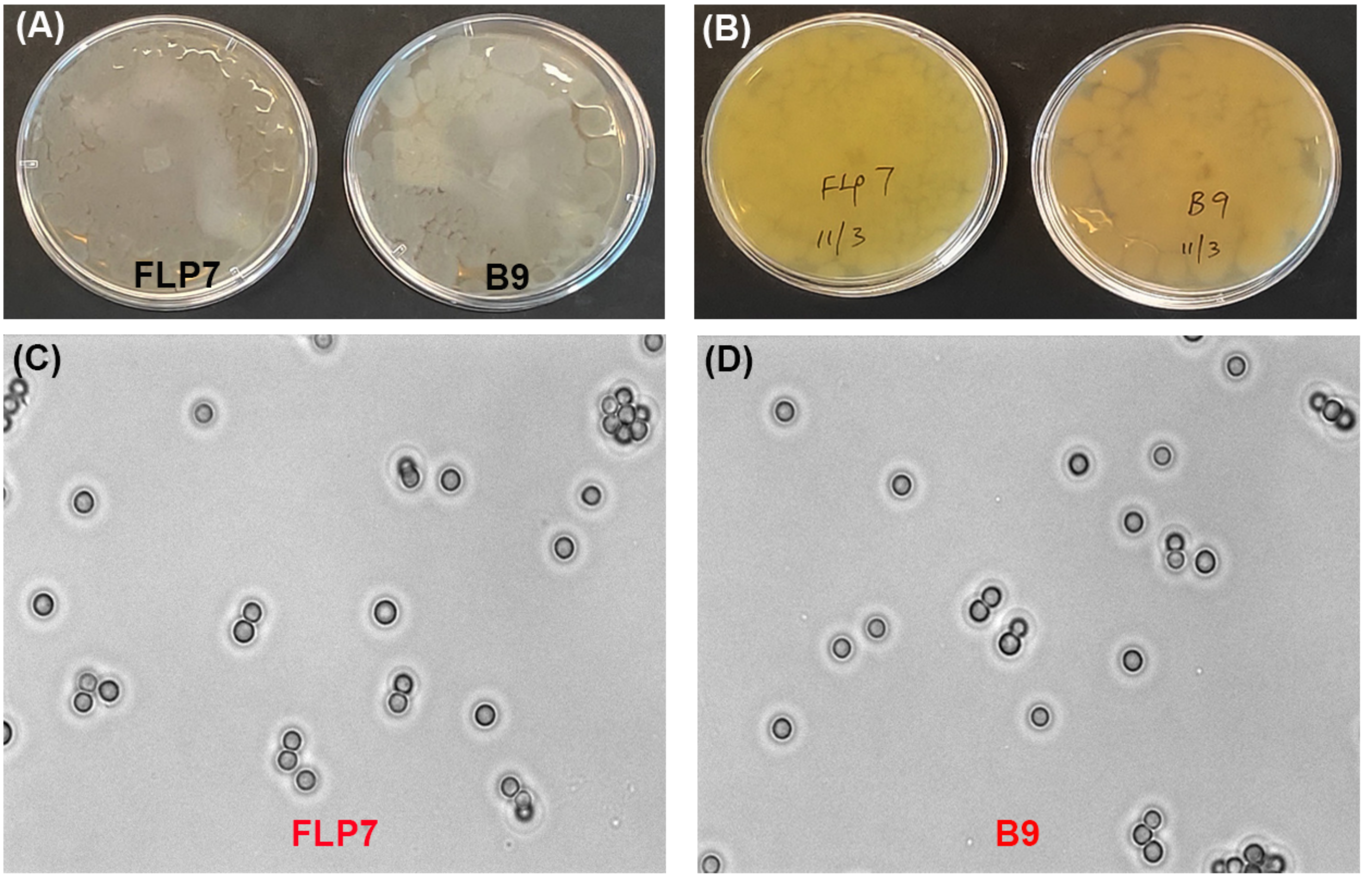

**Supplementary Table S1**

Selected reaction monitoring conditions for protonated or deprotonated forms of the indicated plant hormones ($[M+H]^+$ or $[M-H]^-$)

| Compound Name | Retention Time (RT) | Precursor Ion (Q1) | Product Ion (Q3) | Collision Energy | Polarity |
|---|---|---|---|---|---|
| **Gibberellin GA1** | 2.73 | 346.9 | 272.9 | 32 | Negative |
| | | 346.9 | 228.8 | 30 | |
| | | 346.9 | 145.2 | 30 | |
| **GA3** | 2.67 | 345.1 | 300.9 | 22 | Negative |
| | | 345.1 | 238.8 | 22 | |
| | | 345.1 | 220.8 | 22 | |
| | | 345.1 | 142.9 | 22 | |
| **GA20** | 3.84 | 331.1 | 286.9 | 30 | Negative |
| | | 331.1 | 172.8 | 36 | |
| | | 331.1 | 147 | 30 | |
| **GA4** | 4.84 | 330.9 | 257.2 | 30 | Negative |
| | | 330.9 | 213.2 | 32 | |
| **Auxin / IAA** | 3.13 | 173.8 | 129.8 | 8 | Negative |
| | | 173.8 | 127.9 | 8 | |
| **Cytokinin / Trans-zeatin** | 3.42 | 220.1 | 135.9 | 16 | Positive |
| | | 220.1 | 202.1 | 15 | |
| | | 220.1 | 148.1 | 15 | |
| **Trans-zeatin riboside** | 4.21 | 352.1 | 220.1 | 19 | Positive |
| | | 352.1 | 202.1 | 19 | |

**Supplementary Figure S3**

LC-MS analyses of the indicated phytohormone standards. (**A**) Total ion chromatograms for Gibberellin standards (GA1, GA3, GA4 and GA20) used in this study (**B**) TIC for IAA standard (**C**) TIC for cytokinin standards

(**A**) Total ion chromatogram (TIC) for Gibberellic acid standards (GA1, GA3, GA20, GA4)

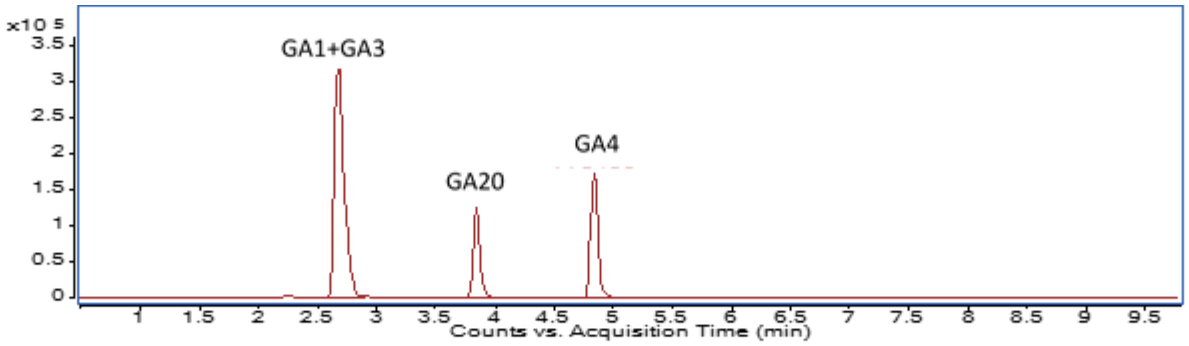


(**B**) TIC for Indole Acetic Acid (IAA/Auxin) standard

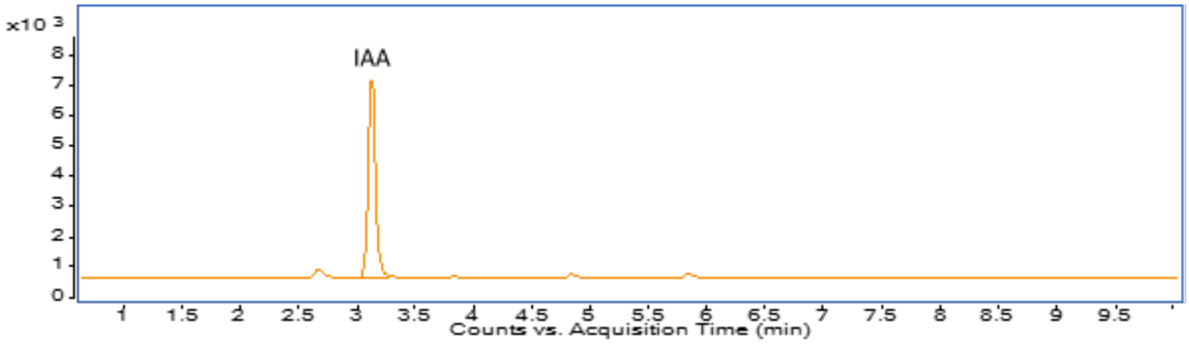


(**C**) TIC for trans-Zeatin and trans-Zeatin-riboside standards

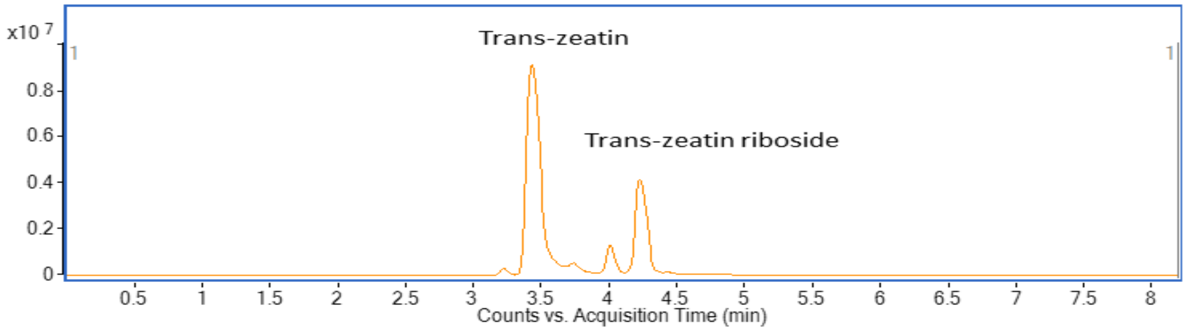

**Supplementary Figure S4 (A-E)**

Multiple Reaction Monitoring transitions for phytohormones showing precursor and product ions **(A)** GA3 **(B)** GA1 **(C)** GA4 **(D)** GA20 **(E)** IAA

**(A)** Multiple Reaction Monitoring (MRM) transitions for GA3 showing precursor ion and fragment ion transitions

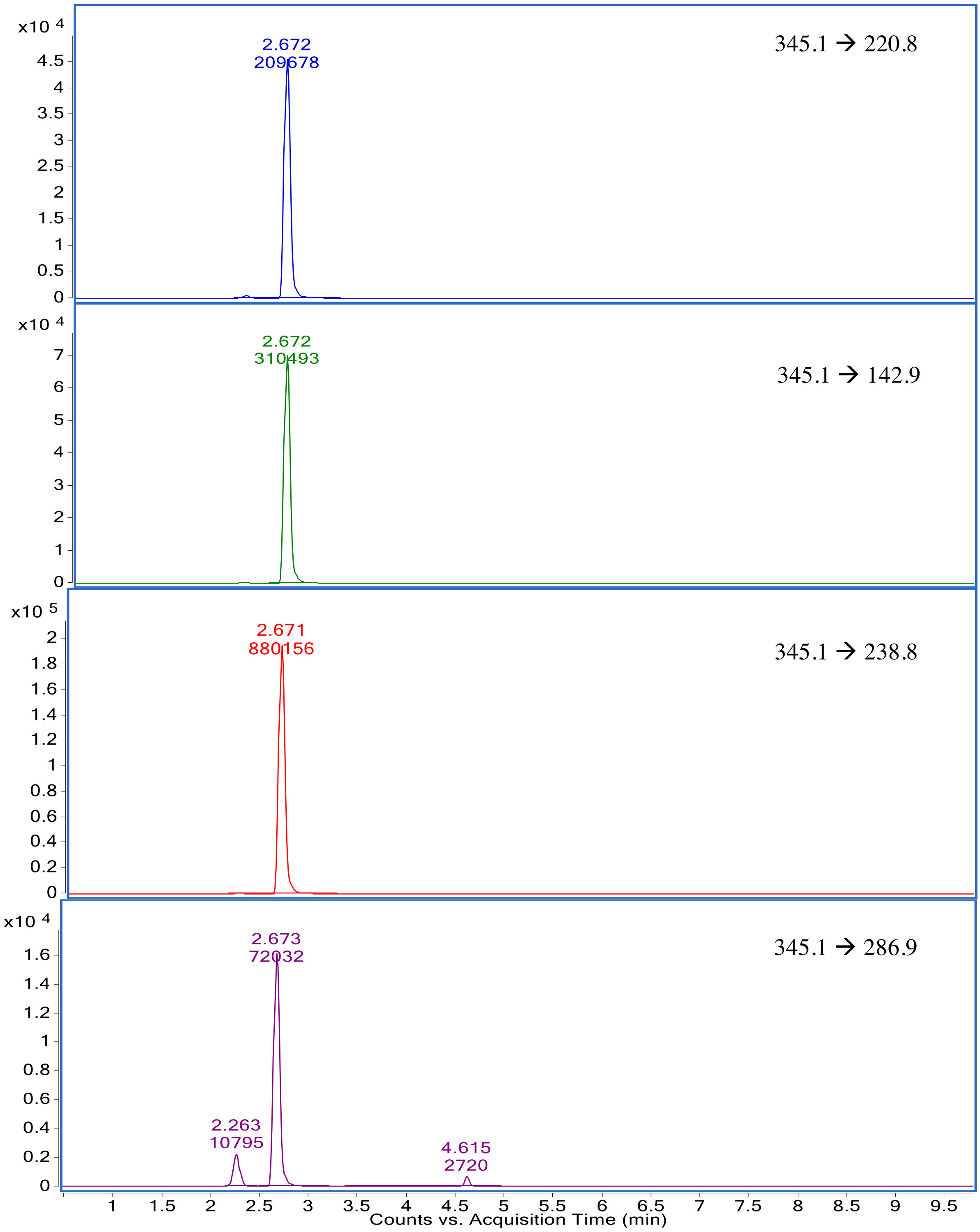

(**B**) MRM transitions for GA1 showing precursor ion and fragment ion transitions

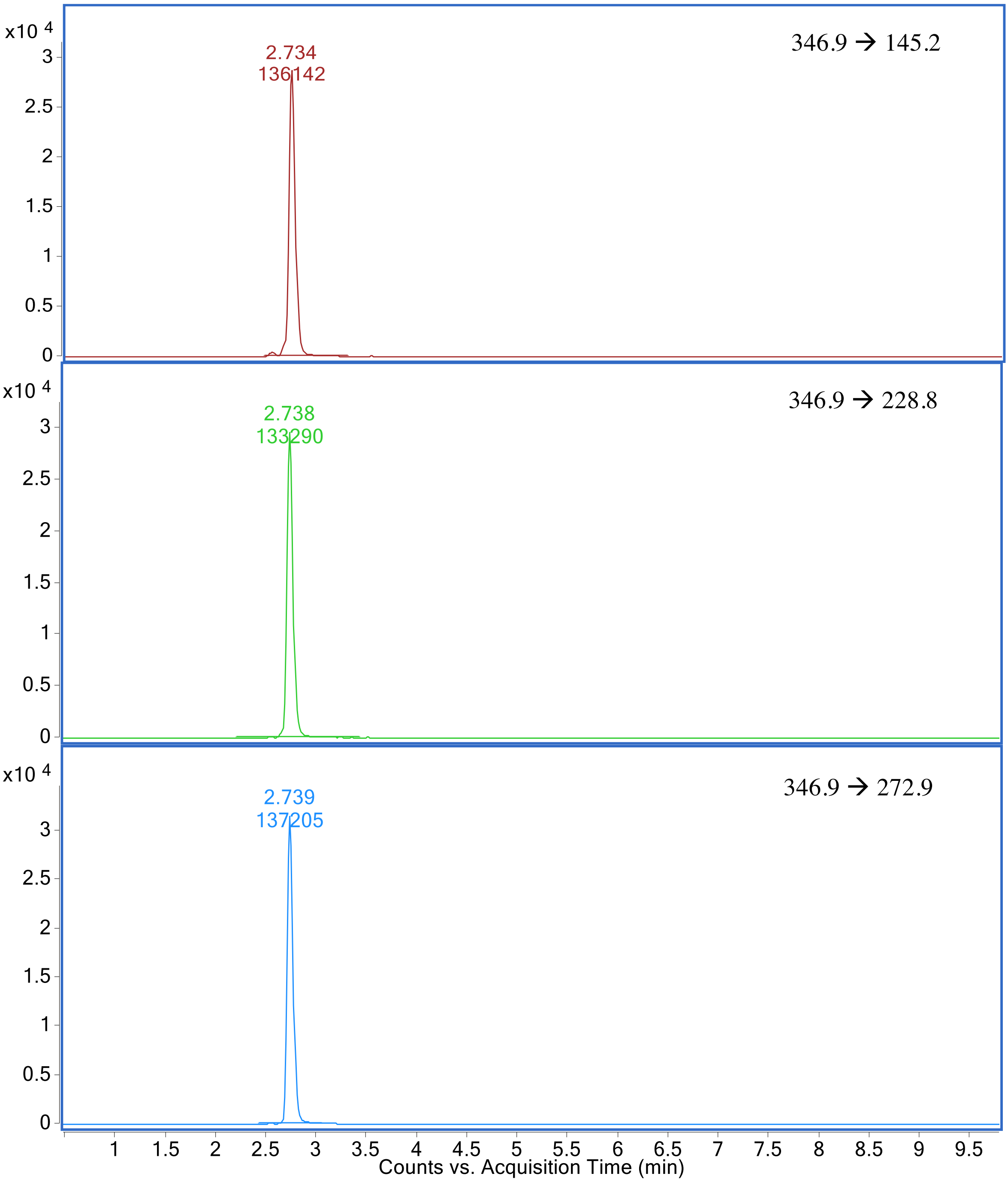

(**C**) MRM transitions for GA4 showing precursor ion and fragment ion transitions

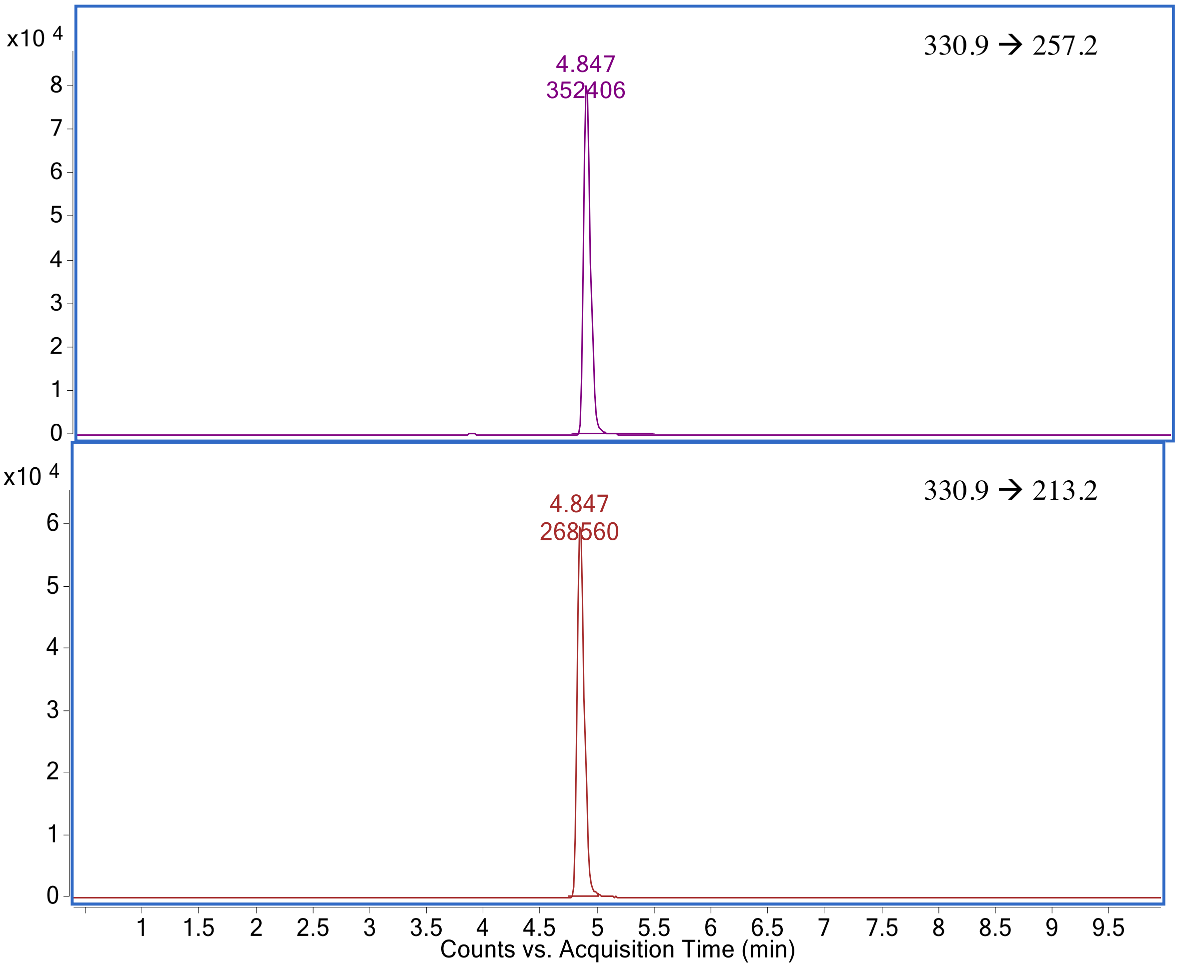

(**D**) MRM transitions for GA20 showing precursor ion and fragment ion transitions

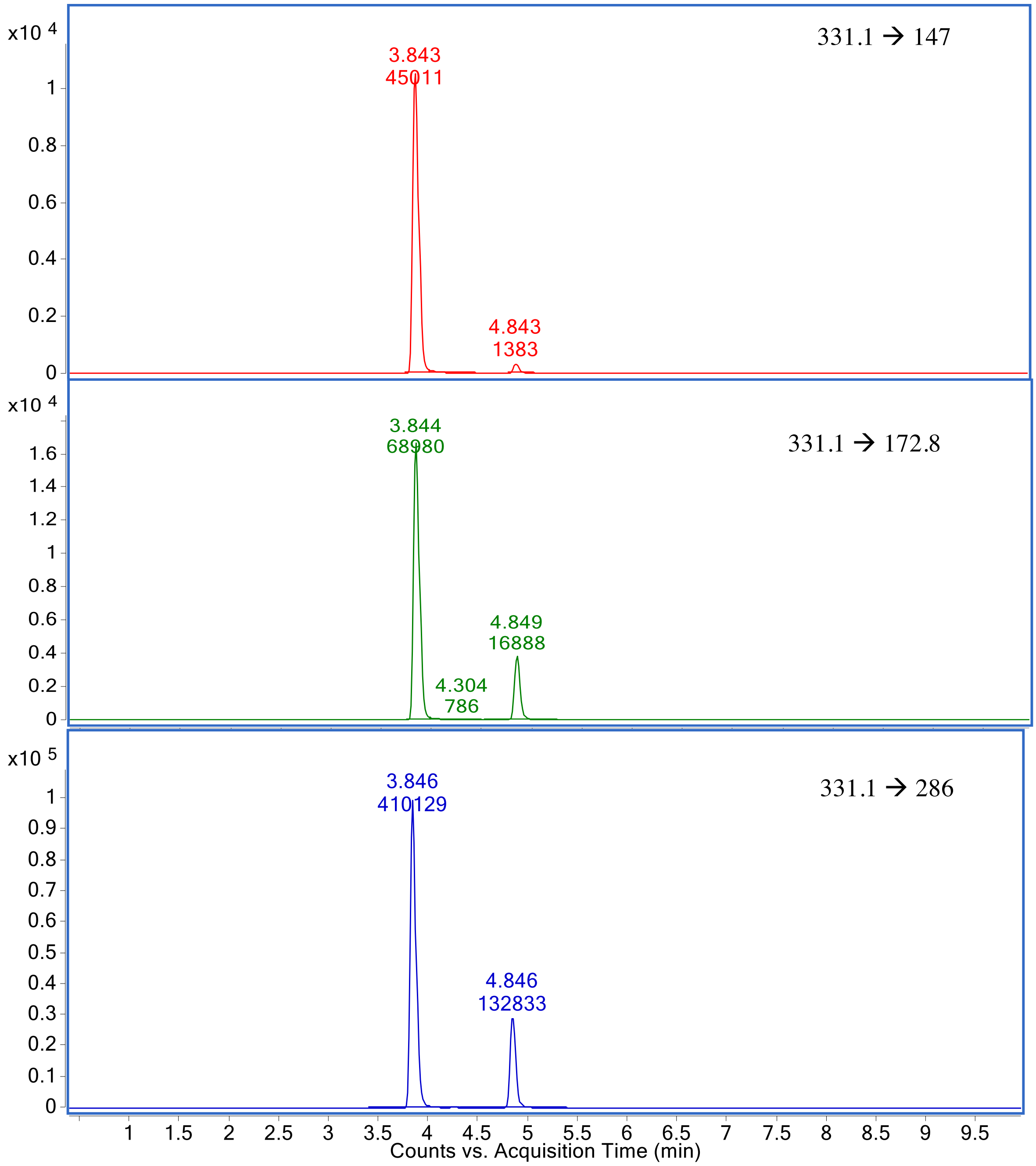

**(E)** MRM transitions for IAA showing precursor ion and fragment ion transitions

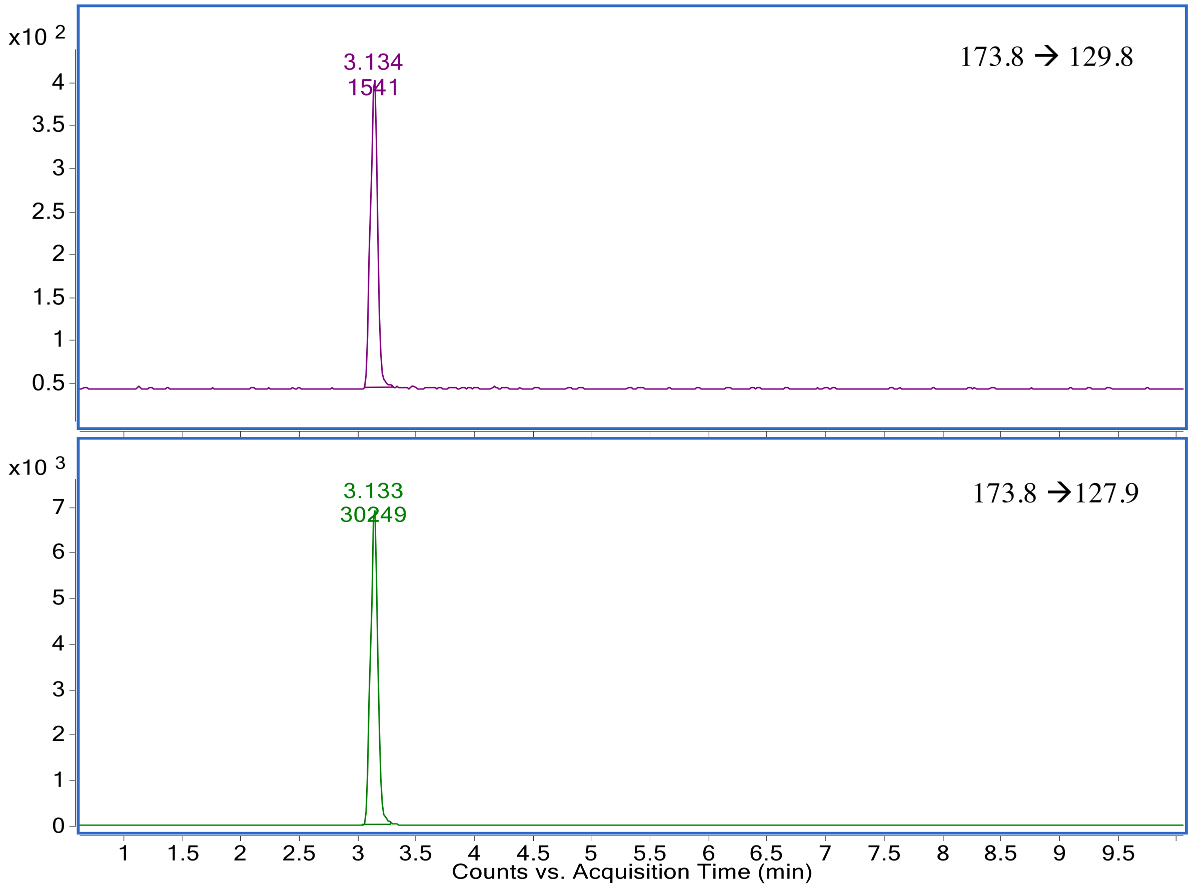

**Supplementary Figure S5:** Multiple Reaction Monitoring transitions for phytohormones showing precursor and product ions (**A**) Trans-zeatin (**B**) Trans-zeatin riboside.

**(A)** MRM transitions for trans-Zeatin-riboside showing precursor ion and fragment ion transitions

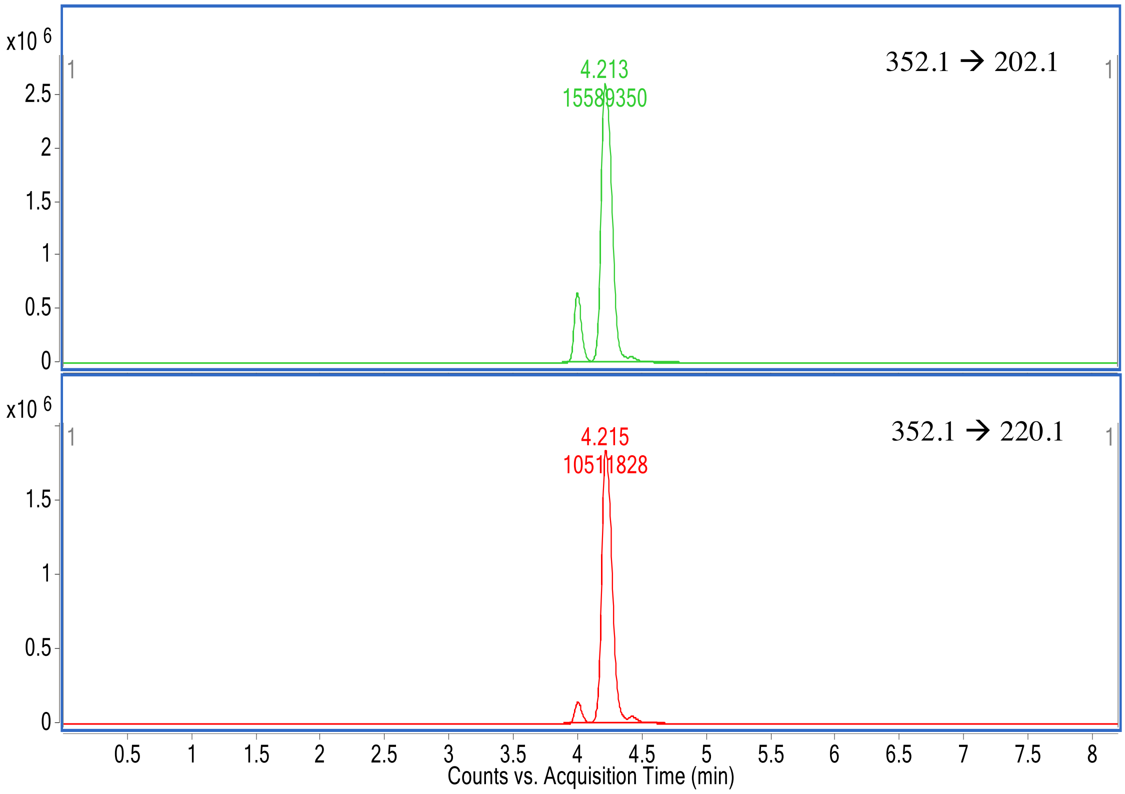


**(B)** MRM transitions for trans-Zeatin showing precursor ion and fragment ion transitions

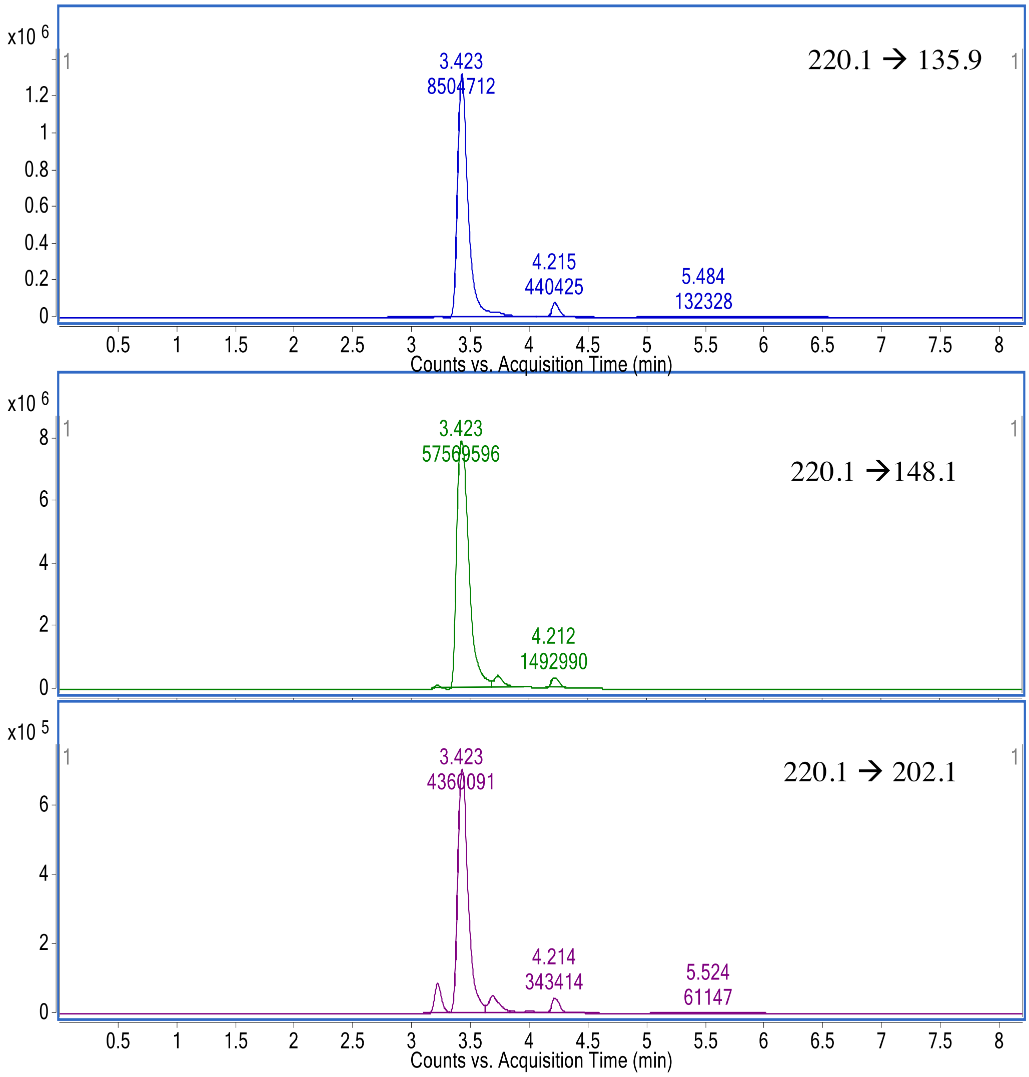

**Supplementary Figure S6**

Phytohormone mimics produced by *P. citrinum*. LC-MS analyses of culture filtrates from B9 and FLP7 isolates of *P. citrinum*. (**A**) Overlay Total ion chromatogram depicting GA1, GA3 and GA20 detected in the FLP7 exudate. (**B**) Overlay TIC between IAA standard and the cell-free culture filtrates of B9 or FLP7 fungal isolates. The presence of IAA in both B9 and FLP7 is evident. The black arrows point to the IAA standard peak, and to the respective IAA peaks in B9 or FLP7. (**C**) Overlay TIC depicting the cytokinin standards (trans-zeatin and trans-zeatin riboside) and the respective detection in cell-free culture filtrates of B9 and FLP-7 fungal isolates along with media extracts (control). Trans-zeatin and trans-zeatin riboside are present in the culture filtrates from both isolates; and in minor amounts in the growth medium (control).

**(A)** Overlay total ion chromatogram (TIC) for two replicates of FLP-7 cell-free culture filtrates showing the presence of GA1+GA3, and GA20.

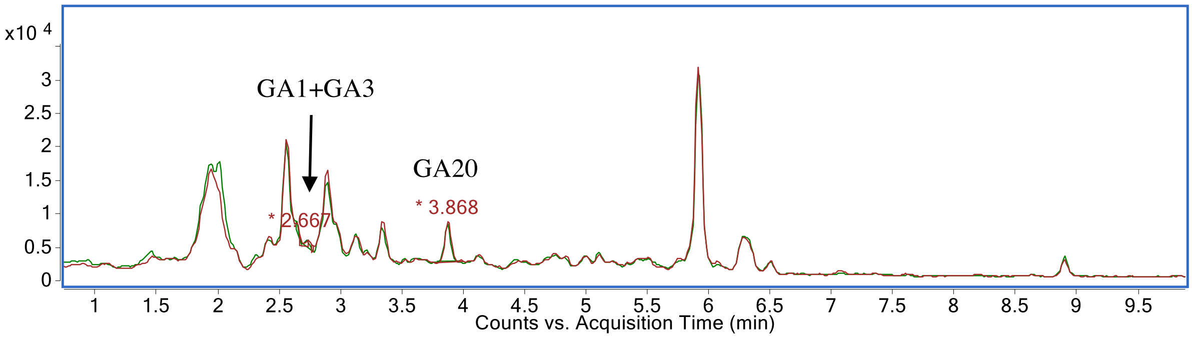


Overlay total ion chromatogram (TIC) between GA20 standard and the cell-free culture filtrates of FLP7 isolate. It shows presence of GA20 in the culture filtrate.

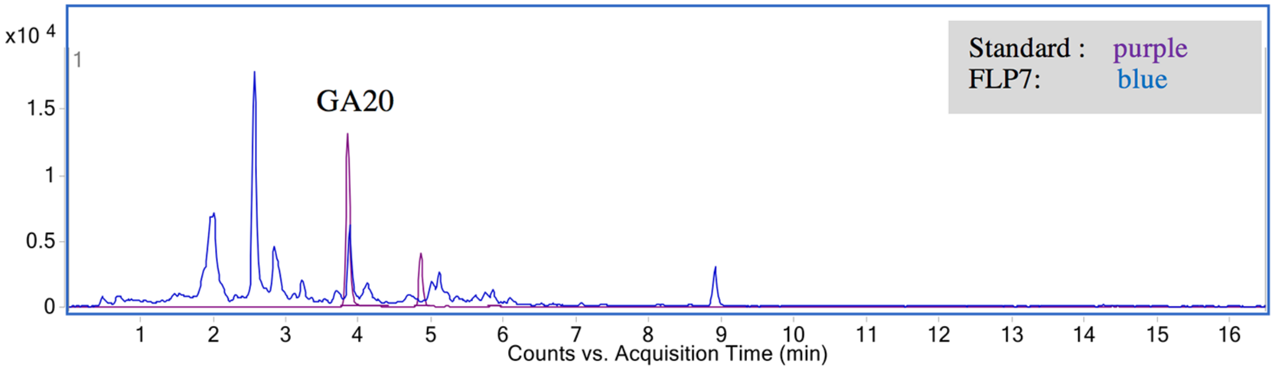


**(B)** Overlay total ion chromatogram (TIC) for IAA in cell-free exudates of the indicated *P. citrinum* isolates (B9 in green, and FLP7 in blue). IAA standard is depicted in red.

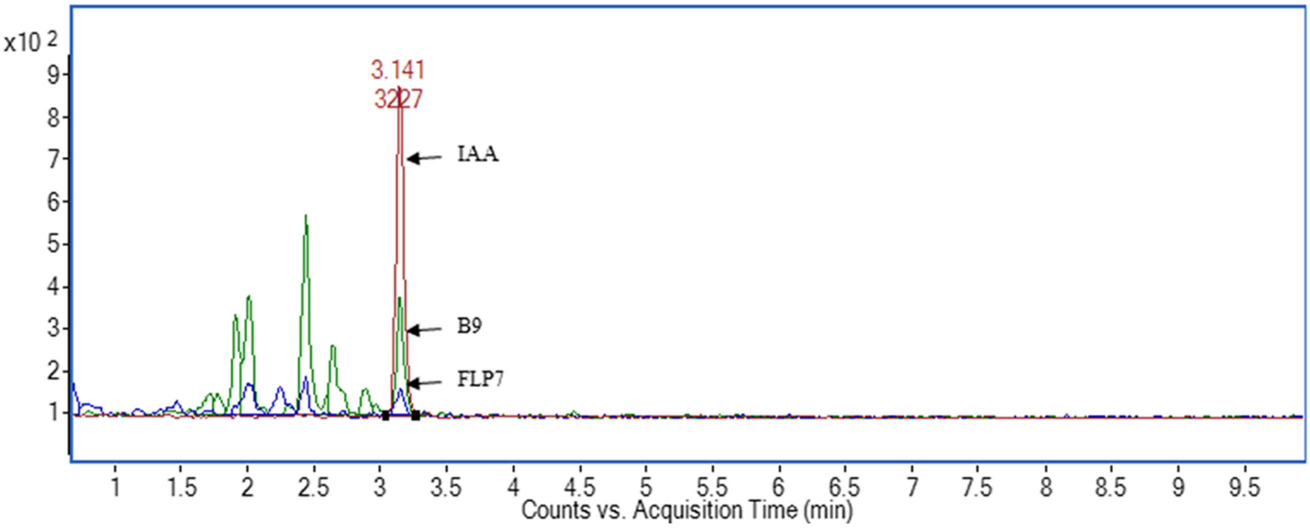

**(C)** Overlay Total Ion Chromatogram (TIC) between trans-Zeatin and trans-Zeatin-riboside standards and cell-free culture filtrates of *P. citrinum* B9 and FLP7 isolates along with culture media extracts (control). Trans-Zeatin and trans-Zeatin-riboside are present in both the filtrates as well as in minor amounts in growth media as well.

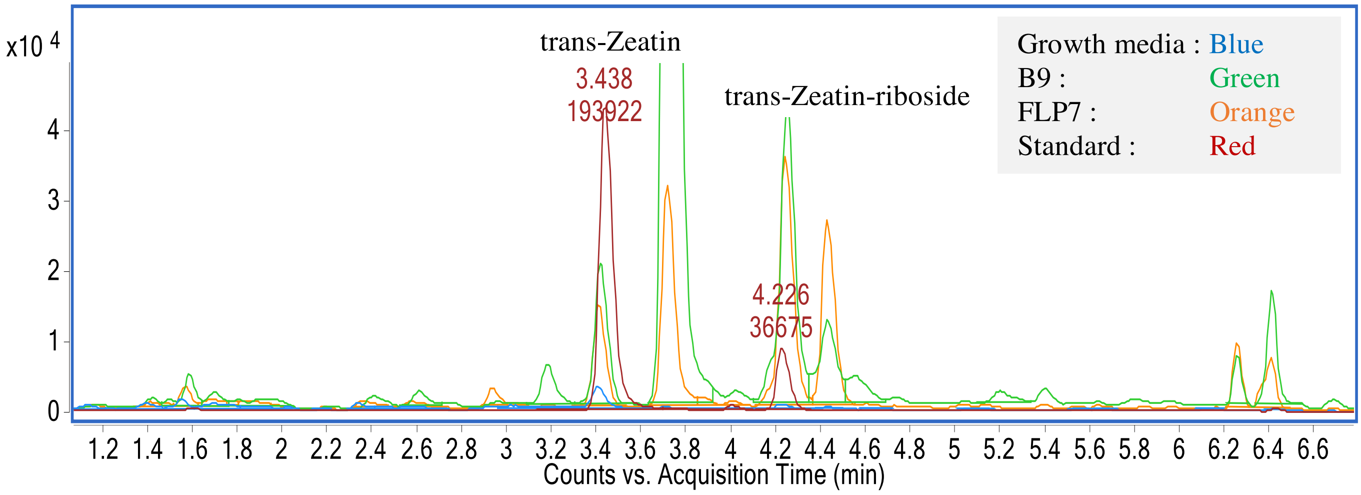
